# A human synovial tendon-on-a-chip models key features of peritendinous adhesions and offers a new approach methodology for testing anti-fibrotic drugs

**DOI:** 10.64898/2026.04.03.716316

**Authors:** Isabelle Linares, Anabelle Creveling, Abdikadir Osman, Nathan Grandwetter, Benjamin L. Miller, James L. McGrath, Hani A. Awad

## Abstract

Peritendinous adhesions are a debilitating complication of tendon injury characterized by excessive matrix deposition and chronic inflammation. Due to limitations of current preclinical models, the underlying mechanisms of adhesion pathogenesis remain poorly defined, and there are no approved drugs to prevent or resolve adhesions. Here, we develop a human synovial tendon-on-a-chip (synToC) that integrates synovial fibroblasts, tendon-resident fibroblasts, immune cells, and vascular endothelium to reconstruct the intrasynovial tendon microenvironment. We show that synovial fibroblast activation promoted tendon contraction and inflammatory cytokine secretion dominated by IL-6, leading to monocyte infiltration and formation of fibronectin- and collagen III-rich matrix bridges between tendon and synovial compartments resembling nascent peritendinous adhesions. These phenotypes emerged even in the absence of exogenous TGF-β1, indicating that synovial fibroblast-mediated crosstalk is sufficient to initiate adhesion-like pathology. Importantly, pharmacological inhibition of the IL-6/JAK/STAT pathway suppressed synovial activation, blunted inflammatory cytokine signaling, and attenuated fibrotic matrix deposition and interfacial adhesion formation. These findings establish the synToC as a human-relevant new approach methodology (NAM) to interrogate the multicellular drivers of tendon adhesions and to accelerate the development of anti-fibrotic therapies.

## Introduction

Adhesions are pathological scar-like connections that form between neighboring tissues and organs following surgery, injury, or inflammation, and occur across multiple anatomical sites, including musculoskeletal, gynecological, cardiovascular, and abdominal tissues^1^. Among these, zone II injuries of the flexor tendon, historically termed “no man’s land”, are particularly challenging, where rates of post-surgical adhesions exceed 40%, and 5% to 15% of patients require secondary surgeries^2, 3^. Current therapeutic strategies, including drugs, physical barrier materials, and physical therapy, often fail to prevent or resolve adhesions and many patients continue to experience impairments in digital range of motion, grip strength, and dexterity^4–6^.

Persistent inflammation and unchecked fibroblast activation have been shown to drive adhesion formation^7^. Activated fibroblasts proliferate and migrate in damaged or healing tissues, promote immune cell recruitment and activation, and secrete cytokines and chemokines that precipitate chronic inflammation and fibrosis^8^. These cells respond to fibrotic signals, including TGF-β1, by activating a myofibroblast phenotype, and deposit excessive extracellular matrix (ECM) components including collagen and fibronectin, forming scar tissue that ultimately compromises the regeneration of native tendon structure and normal function. Despite decades of investigation, the cellular origins of the fibroblasts responsible for adhesions remain a source of debate^9^. Evidence from animal models supports the hypothesis that tendon adhesions arise from an imbalance between the intrinsic (tendon-derived) and extrinsic (synovial sheath and surrounding tissue-derived) healing responses^10^. Although both intrinsic tendon cells and synovial sheath cells upregulate TGF-β1 pathways early after injury^11^, sheath cells show greater reactivity, migratory capacity, and matrix synthesis in response to TGF-β1^9, 12–14^. Fibroblast-like synoviocytes (FLS), which constitute the major cellular population of the tendon sheath, therefore represent strong candidates as key effector cells linking extrinsic healing to pathological tendon fibrosis and adhesions.

While the role of FLS in peritendinous adhesions remains poorly defined, their contribution to synovial inflammation and fibrosis is well established in other musculoskeletal pathologies. In rheumatoid arthritis (RA), FLS sustain synovial inflammation through direct cell-cell interactions and secretion of soluble mediators^15^. FLS have also been implicated in synovial fibrosis in osteoarthritis (OA), exhibiting increased proliferation and secretion of pro-inflammatory mediators, including IL-6 and IL-8^16^, and upregulating TGF-β1-associated genes linked to FLS activation (ACTA2, CDH11) and matrix synthesis (COL1A1, COL3A1, THBS1, FN1), supporting their direct involvement in synovial fibrosis^16, 17^. Furthermore, the crosstalk between FLS and other synovial cells is critical. Recent evidence from refractory RA synovial biopsies identified a complex synovial microenvironment in which inflamed vascular endothelial cells drive FLS activation by creating a spatial gradient of fibrogenic TGF-β1 signaling in the synovium, which contributes to treatment resistance^18^. Interestingly, while tendons are mostly avascular, the scar tissue constituting peritendinous adhesions is highly vascularized. Consequently, a similar crosstalk between inflamed endothelial cells, leukocytes, and fibroblasts likely facilitates immune cell trafficking and orchestrates myofibroblast microenvironment (MME) activation in peritendinous adhesions^19^. We posit that multicellular signaling within the inflammatory MME could be key to identifying therapeutic targets for the resolution of adhesions.

While animal models are widely used to study intrasynovial flexor tendon healing, fundamental limitations restrict their utility to elucidate the multicellular mechanisms that drive adhesion formation. Rodent models provide powerful genetic tools and are well suited for tracing specific cell populations.^20^ However the immune systems and tendon architecture differ between humans and rodents, creating a mismatch in response to drug treatments.^21^ Large animal models improve replication of surgical and clinical factors but are costly, low-throughput, and poorly suited for systematic manipulation or longitudinal analysis of specific cell-cell signaling pathways. As emphasized by recent consensus efforts, no existing animal model fully captures the spatial, temporal, and cellular complexity of tendon adhesion pathogenesis^22^. Furthermore, current *in vitro* systems do not adequately recapitulate the dynamic intrasynovial tendon microenvironment, lacking key features such as vascular-stromal interfaces, immune cell trafficking, mechanical cues, and spatiotemporally regulated paracrine signaling^23^. Consequently, these reductionist models cannot adequately simulate the multicellular interactions in fibrotic adhesions, and research must prioritize new approach methodologies (NAMs), particularly tissue- and organ-on-chip platforms, to provide physiologically-relevant disease models. By generating more accurate and predictive human-specific data, these models promise to accelerate and de-risk drug development while complementing traditional animal models.

We previously engineered a human tendon-on-a-chip (hToC) featuring a 3D tendon construct interfaced with a perfused vascular channel across an ultrathin nanomembrane with dual (micro/nano) porosity enabling controlled immune cell trafficking and recapitulation of inflammatory and fibrotic features of tendon pathology driven by TGF-β1^19, 24^. This work established a human-relevant model of intrinsic tendon fibrosis arising from endothelial-immune-fibroblast crosstalk within the MME. However, like other existing tissue-on-chip fibrosis models, the hToC does not capture the contribution of extrinsic synovial sheath cells, which are strongly implicated in peritendinous adhesion formation.

Here, we report the development of an intrasynovial tendon-on-a-chip, which captures physiologically and clinically relevant features of peritendinous adhesion. This is accomplished by incorporating a vascularized synovial lining into the hToC platform to create a human synovial tendon-on-a-chip (synToC) that reconstitutes inflammatory vascular-synovial-tendon crosstalk and enables formation and quantification of adhesion-like ECM structures. We investigated the hypothesis that the FLS-seeded synovial hydrogel lining drives tendon adhesion formation by amplifying inflammatory signaling and promoting immune cell recruitment and fibrotic matrix deposition. We demonstrate that FLS are sufficient, in the absence of exogenous TGF-β1 treatment, to induce cellular, molecular, and structural features of peritendinous adhesions, including enhanced monocyte transendothelial migration, synovial hyperplasia, and deposition of fibronectin- and collagen III-rich bridging ECM that physically connects the synovial and tendon compartments. Importantly, the model enables rational testing of drugs based on upregulated cytokines, such as IL-6, and quantification of functional improvements in synovial hyperplasia, inflammatory signaling, and tissue-scale adhesion metrics that cannot be assessed in existing fibrosis models or conventional in vitro assays. Considering that clinically used imaging and range-of-motion metrics are directly associated with these physical adhesions^25–27^, the synToC provides a first-in-kind, clinically relevant new approach methodology (NAM) to dissect intrinsic and extrinsic drivers of tendon adhesions and to perform preclinical, mechanism-informed evaluation of anti-adhesion therapies.

## Results

### Incorporating a vascularized-synovial lining to model the intrasynovial tendon microenvironment in the synToC

As with the hToC^19, 24^, a key feature of the synToC platform is its modular design, which enables independent culture of the tendon and vascular-synovial compartments prior to integration (**Fig. 1a**). To recapitulate the intrasynovial tendon microenvironment in the bottom compartment, primary human tendon fibroblasts and monocytes were encapsulated in a type I collagen hydrogel and cultured with macrophage colony-stimulating factor (M-CSF) for 6 days (Day-6 to Day 0) to promote monocyte to macrophage differentiation and reconstitute the early post-injury tendon inflammatory niche. For the vascular-synovial compartment, a type I collagen hydrogel embedded with or without primary fibroblast-like synoviocytes (±FLS) was polymerized on the basal surface of the porous nanomembrane. Twenty four hours later, ECs were seeded on the opposite apical side of the nanomembrane and cultured for an additional 24 hours to establish a robust vascular-synovial interface. At Day 0, the vascular-synovial and tendon components were assembled into the complete synToC, monocytes were introduced to the vascular compartment, and serum-free media with or without 10 ng ml^-1^ TGF-β1 was added to the bottom (tendon) compartment. CellTracker labeling demonstrated compartmental localization of FLS and tendon fibroblasts within their respective regions and demonstrated transendothelial migration of monocytes to the synovial hydrogel and underlying tendon tissue (**Fig. 1b**).

**Figure 1.**
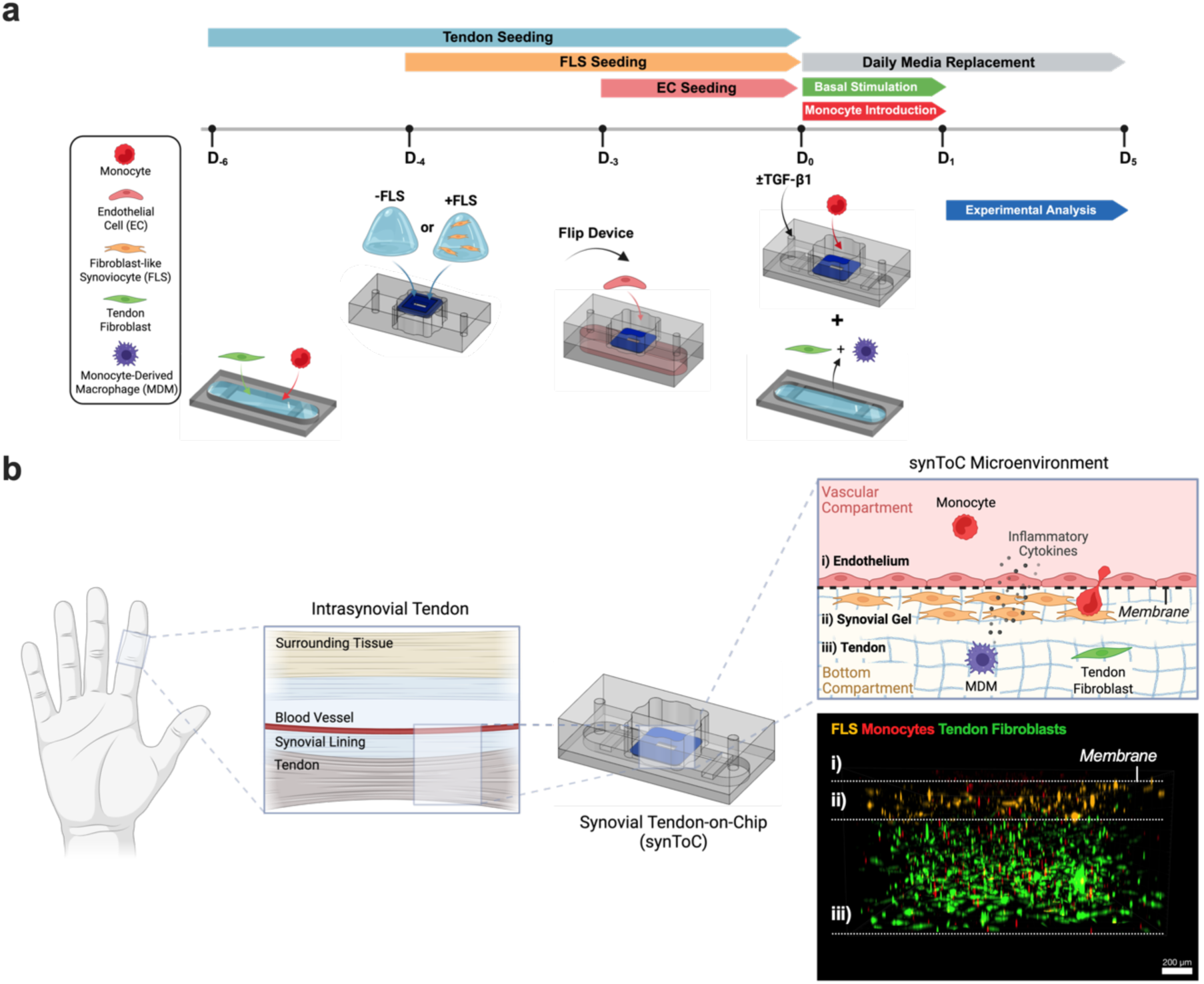
Reconstitution of the intrasynovial tendon microenvironment in the synToC platform. **a**, Culture timeline outlining the steps to incorporate a synovial layer. A tendon construct containing primary tendon fibroblasts and macrophages was matured in parallel in the bottom chamber. FLS are encapsulated in a type I collagen hydrogel and added to the basal side of the porous nanomembrane (acellular collagen for - FLS), followed by seeding vascular endothelial cells on the apical side to form the vascularized synovial compartment. At Day 0, the vascular-synovial compartment is joined with the bottom tendon compartment. Freshly isolated monocytes are added to the top compartment with or without TGF-β1 in the bottom compartment. Experimental analyses are performed from Day 1 to Day 5. **b,** Schematic detailing the functional unit of an intrasynovial tendon that is recapitulated in the synToC. The confocal immunofluorescence image corresponds to the cross-sectional view of representative cell types within the device (i) endothelium, ii) synovial gel, and iii) tendon) and demonstrates the spatial localization of FLS (orange), monocytes (red), and tendon fibroblasts (green) at Day 1 of culture. ECs and tendon resident monocyte-derived macrophages (MDM) are present but not labeled.

To model the vascular synovial microenvironment within the synToC, we established a spatially organized fibroblast-like synoviocyte (FLS)-endothelial cell (EC) co-culture, in which ECs form a monolayer to model a vascular wall on the apical nanomembrane surface and an FLS-seeded collagen hydrogel is polymerized on the basal side. First, we evaluated different culture media conditions and identified that a 1:1 mix of synoviocyte growth media and endothelial growth media was optimal to simultaneously preserve endothelial junction integrity (VE-cadherin) and synovial fibroblast phenotype (cadherin-11) when these cells are cultured separately in monolayer **(Supplementary Fig. S1).**

To mimic the vascularized synovial sheath that surrounds intrasynovial tendons^28, 29^, we encapsulated FLS within a three-dimensional type I collagen hydrogel. The hydrogel was then polymerized on the basal side of the porous nanomembrane and cultured for 24 h before flipping the device and seeding ECs on the apical side (**Fig. 2a**). Using immunofluorescence staining, we verified the expression of VE-cadherin endothelial junctions across the vascular monolayer and maintenance of FLS expression of cadherin-11 (**Fig. 2b**), with discernable compartmentalization across the membrane. We also confirmed the expression of PRG4, a marker of synovial cells that line tendon sheaths providing lubrication and cytoprotection^30^, in the FLS-laden synovial hydrogel in close proximity to the endothelial barrier (**Fig. 2c**). To assess the co-culture response to a fibrotic stimulus, TGF-β1 (10 ng ml⁻¹) was introduced to the basal compartment at Day 3. Fibrosis-associated collagen III expression remained stable between Day 3 and Day 6 without TGF-β1 stimulation, but significantly increased at Day 6 with TGF-β1 treatment (**Fig. 2d**). Ki67 immunostaining also demonstrated a significant increase in a proliferative FLS phenotype at Day 6 with and without TGF-β1 treatment (**Fig. 2e**). Total FLS cell density increased by Day 6 regardless of TGF-β1 stimulation (**Fig. 2f**).

**Figure 2.**
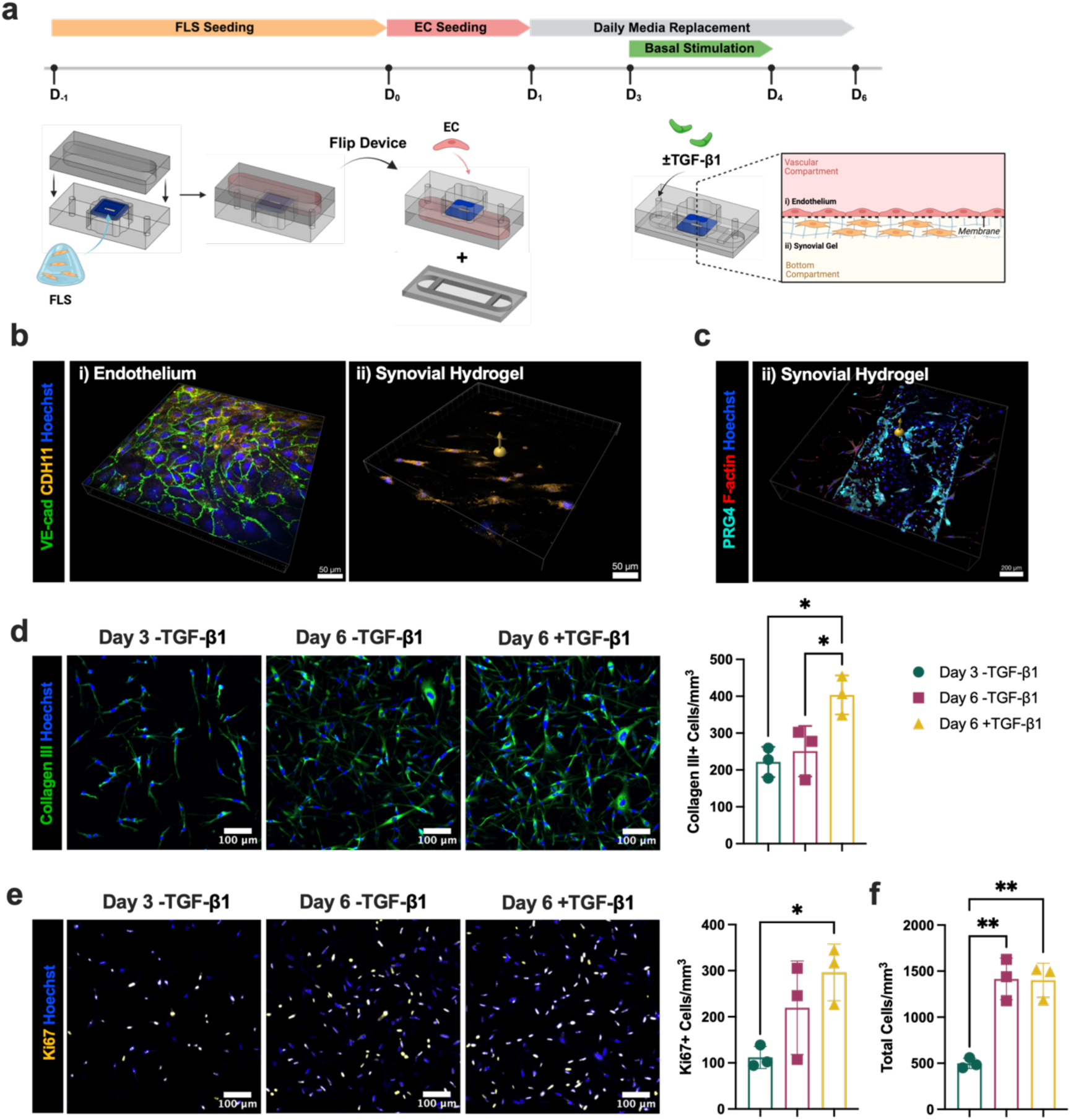
Establishing and validating a vascularized synovial co-culture in the synToC. **a**, An overview of the culture timeline to establish an FLS-EC co-culture. FLS are encapsulated in a type I collagen hydrogel and added to the basal side of the nanomembrane. After 24 h, ECs are added to the top vascular compartment and form a vascular barrier over 24 h. Media is replaced daily until Day 3 (D_3_) at which point an inflammatory stimulus (10 ng ml⁻¹ TGF-β1) is added to the basal compartment. **b,** FLS-EC co-culture at Day 3 stained for VE-cadherin (green), cadherin-11 (yellow), and Hoechst (blue). Maximum intensity projection taken at 10x (left). i) z-slice focusing on the endothelial monolayer. ii) z-slice focusing near the bottom of the FLS gel, demonstrating robust VE-cadherin junctions in the ECs above the FLS gel and compartmentalization of cadherin-11 expression in the FLS gel. **c,** Synovial hydrogel stained for PRG4 (cyan) and F-actin (red) showing a z-slice in the FLS gel focusing just under the endothelium layer on the nanomembrane demonstrating robust PRG4 secretion from FLS closest to the endothelial barrier. **d,** Immunofluorescence images of collagen III (green) expression in the synovial gel, with Hoechst (blue) as nuclear counterstain. Density of collagen III+ cells upon stimulation with TGF-β1 on Day 6. **e,** Immunofluorescence images of nuclear Ki67 (yellow) expression in the synovial gel, with Hoechst (blue) as nuclear counterstain. The number of Ki67+ cells mm^-3^ significantly increased at Day 6 with TGF-β1 stimulation. **f,** Quantification of the total cell density in the synovial gel, showing comparable increases in cell counts at Day 6 regardless of TGF-β1 stimulation. One-way ANOVA with Tukey’s post-hoc test: n=3 devices per condition, *p<0.05, **p<0.01.

### Synovial FLS are sufficient to induce intrinsic tendon fibrotic phenotypes

We next investigated whether incorporating an acellular or FLS-laden synovial hydrogel, with or without supplementation with TGF-β1, modulates fibrotic remodeling within the tendon hydrogel. TGF-β1 is an important growth factor that contributes to pathological fibrosis and tendon adhesions^11^. Macrophages, fibroblasts, and endothelial cells secrete TGF-β1 at the site of injury, which promotes fibroblast to myofibroblast differentiation and proliferation^31^. α-SMA^+^ myofibroblasts are an established marker for fibrosis in injured tendons^31–33^. In the absence of TGF-β1 supplementation, tendon hydrogel contraction was significantly increased in devices with FLS-laden synovial gels (+FLS) compared to acellular synovial gels (-FLS) as early as Day 3, with increased differences by Day 5 (**Fig. 3a**). Upon supplementation with TGF-β1, +FLS and -FLS devices exhibited comparable bulk tissue contraction throughout the duration of culture (**Fig. 3b**). Notably, +FLS devices exhibited significantly greater maximum contraction at Day 5 in the absence of TGF-β1 (**Fig. 3c**). Maximum tendon contraction at Day 5 in +FLS devices under -TGF-β1 conditions was consistent with increased collagen packing density, as evidenced by significant increases in second harmonic generation (SHG) signal intensity in the tendon construct relative to -FLS devices **(Supplementary Fig. S2)**.

**Figure 3.**
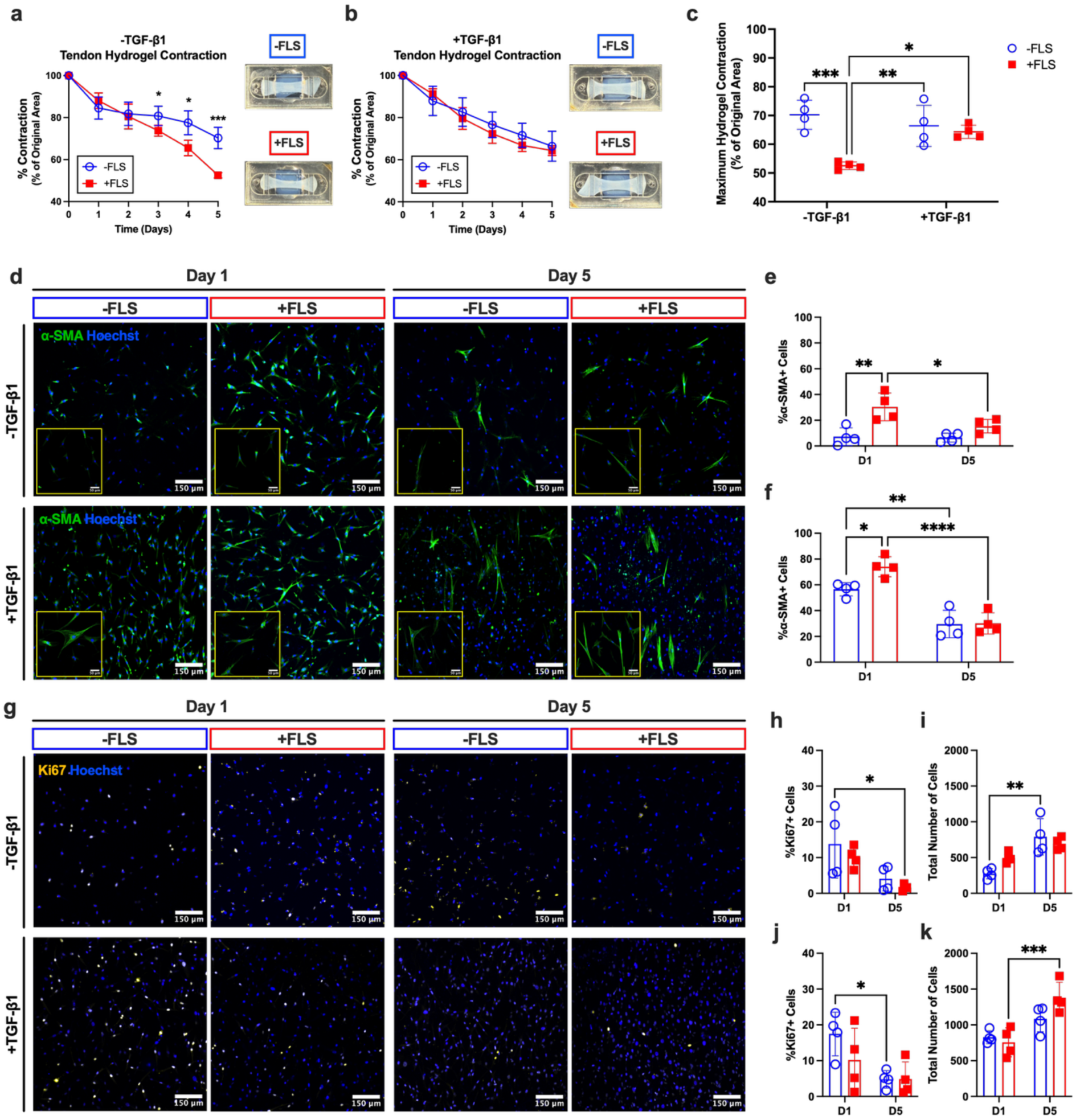
Incorporating FLS in a synovial lining is sufficient to induce significant tendon hydrogel contraction and results in increased levels of α-SMA in the tendon hydrogel even in the absence of TGF-β1. Tendon hydrogel contraction plotted as the percent of the original hydrogel area in devices **a,** without TGF-β1 and **b,** with TGF-β1 stimulation. Representative images of tendon hydrogels in a and b are shown at Day 5. Data are mean ± s.d. (n=4 devices per condition); Bonferonni corrected t-tests at each time point: n=4 devices, *p<0.05, ***p<0.001. **c,** Quantification of maximum hydrogel contraction. Two-way ANOVA with Tukey’s post-hoc test; *p<0.05, **p<0.01, ***p<0.001. **d,** α-SMA (green) immunofluorescence in the tendon hydrogel comparing devices ±FLS, with or without exogenous TGF-β1. Quantification of α-SMA expression in devices **e,** without TGF-β1 and **f,** with TGF-β1 stimulation, reported as the percent positive expressing cells. **g,** Nuclear Ki67 (yellow) immunofluorescence in the tendon hydrogel comparing devices ±FLS with or without exogenous TGF-β1. Quantification of Ki67 expression in devices **h,** without TGF-β1 and **j,** with exogenous TGF-β1. **i, k,** Quantification of the total number of cells per field of view in the tendon hydrogel. Data are mean ± s.d. (n=4 devices per condition); two-way ANOVA with Tukey’s post-hoc test; *p<0.05, **p<0.01, ***p<0.001, ****p<0.0001.

Analysis of immunofluorescence staining of α-SMA in the tendon construct demonstrated that, under basal conditions (-TGF-β1), +FLS devices exhibited a significantly higher proportion of α-SMA⁺ cells at Day 1 compared to -FLS controls (**Fig. 3d**), indicating that FLS induce early myofibroblast differentiation in the tendon hydrogel. Upon supplementing with TGF-β1, we observed additive increases in α-SMA expression at Day 1 for both -FLS and +FLS devices (**Fig. 3e,f**), with 74.1±7.8% α-SMA⁺ tendon cells in the +FLS,+TGF-β1 devices compared to 30.41±10.7% in the +FLS,-TGF-β1 devices. Despite an overall decrease in the number of α-SMA^+^ cells over time across all conditions, high magnification images depict the formation of mature stress fibers at Day 5 compared to diffuse α-SMA expression in the tendon fibroblasts at Day 1 (**Fig. 3d**). Furthermore, the percentage of α-SMA+ cells was significantly elevated at Day 1 in +FLS devices compared to -FLS devices, regardless of TGF-β1 treatment (**Fig. 3e,f)**. There were no significant FLS- or TGF-β1-dependent expression differences in Ki67, a nuclear marker of cellular proliferation that has been observed in highly proliferative cells in adhesive peritendinous tissues^34, 35^ (**Fig. 3g**). While the percentage of Ki67^+^ cells tended to decrease over time in all conditions (**Fig. 3h,j**), cell counts were elevated at Day 5 (**Fig. 3i,k**) suggesting that the enhanced contraction and myofibroblast differentiation were driven by proliferative phenotypes at Day 1.

### FLS induce synovial hyperplasia and monocyte infiltration

To further investigate the FLS-laden synovial gel inflammatory response in the absence of exogenous TGF-β1, we evaluated proliferative activity, reparative fibroblast markers, inflammatory synovial markers, and immune cell recruitment over time. Immunofluorescence staining of nuclear Ki67 as a proliferation index demonstrated elevated levels of cellular proliferation within the synovial gel at Day 1. Although Ki67⁺ cell density declined by Day 5, the total synovial density, as quantified from nuclei counts, increased significantly over time (Day 1: 1182±121.7 cells mm^-3^; Day 5: 1680±301.8 cells mm^-3^), indicating net cellular expansion despite the decline in proliferative index between days 1 and 5 (**Fig. 4a**).

**Figure 4.**
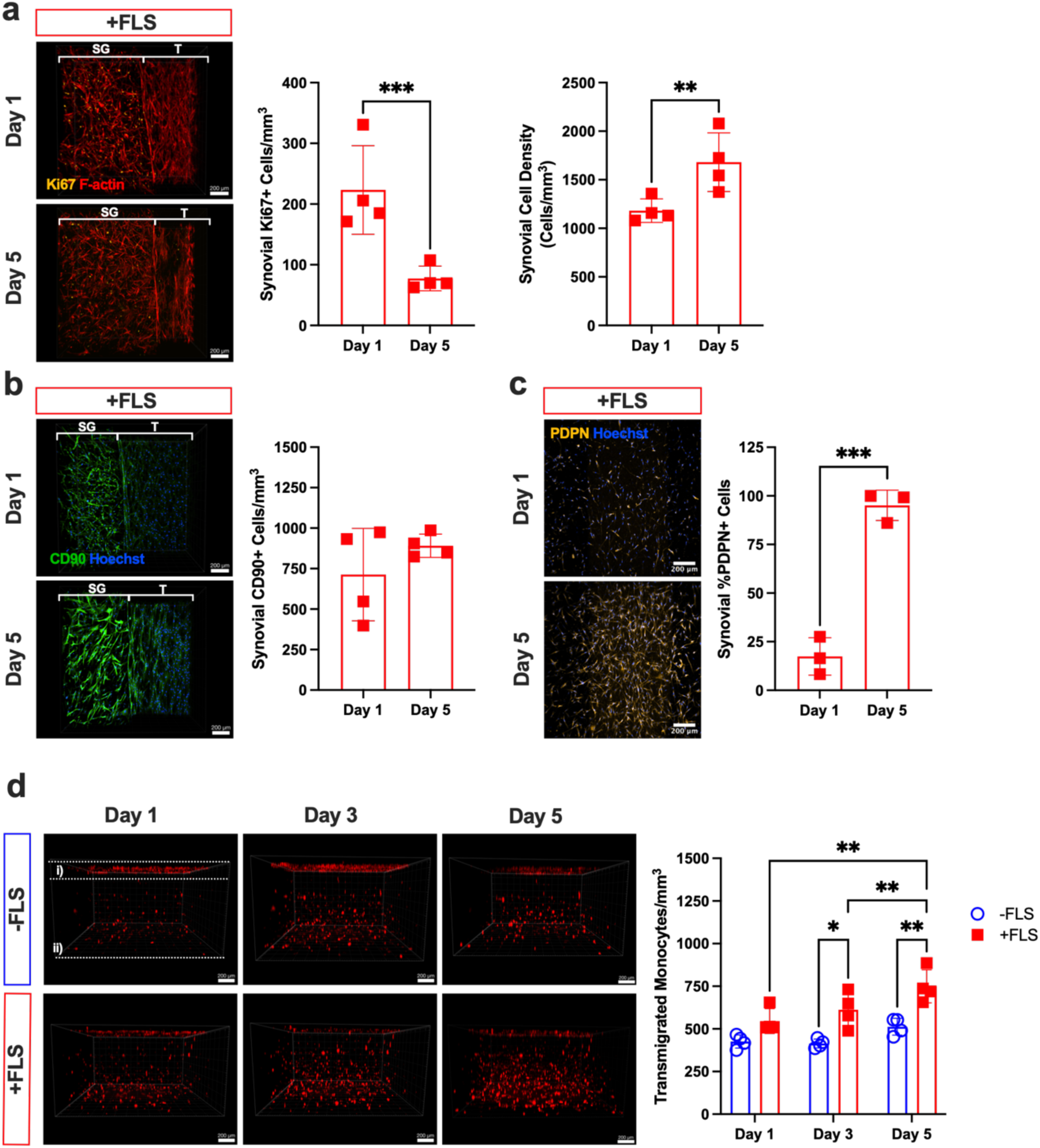
FLS induce synovial hyperplasia, inflammatory activation, and monocyte infiltration under basal conditions. **a**, Immunofluorescent confocal images of Ki67 (yellow) in the synovial gel (SG) adjacent to the tendon gel (T) at Days 1 and 5 without exogenous TGF-β1. Quantification of Ki67+ cells and total synovial cell density demonstrate elevated proliferation at Day 1 and an increased overall cell density in the synovial gel by Day 5. **b,** Immunofluorescent confocal images and quantification of CD90 (green) expression with Hoechst (blue) as nuclear counterstain at Days 1 and 5 demonstrate enrichment of reparative CD90+ cells in the synovial gel. **c,** Immunofluorescent confocal images and quantification of podoplanin (PDPN, yellow) expression within the synovial gel, demonstrating significant upregulation from Day 1 to Day 5. **d,** Three-dimensional confocal imaging of live CellTracker-labeled monocytes (red) at Days 1, 3, and 5 showing transendothelial migration into the (i) synovial and (ii) tendon compartments (±FLS). Quantification of monocyte infiltration, reported as the number of transmigrated monocytes mm^-3^, shows time-dependent increase in monocyte infiltration in +FLS compared to -FLS devices at Days 3 and 5. Data are mean ± s.d.(n=4 devices per condition); Two-way ANOVA with Tukey’s post-hoc test; *p<0.05, ***p<0.001.

We next assessed CD90, a cell surface glycoprotein that regulates fibrosis and has also been established as a marker of reparative fibroblasts^36, 37^. We observed robust expression of CD90 in the FLS-laden synovial hydrogel at both Day 1 and Day 5 (**Fig. 4b**), consistent with enrichment of a fibroblast subset linked to adhesions^1^. We also observed a significant increase in podoplanin (PDPN) expression within the synovial gel from Day 1 to Day 5 (**Fig. 4c**). PDPN is a marker of inflammation in stromal fibroblasts in tendinopathy^38^, tenosynovitis^39^, and tendon adhesions^40^. At Day 1, PDPN expression was detected in a sparse subset of synovial cells, whereas by Day 5 PDPN signal intensity was broadly distributed throughout the synovial gel, along with increased FLS density and morphological elongation.

In contrast, the acellular synovial hydrogel contained a sparse population of fibroblast-like cells that seem to have migrated outward from the tendon hydrogel rather than expansion of a resident synovial population observed in the +FLS synovial gels. CD90 expression within the - FLS synovial compartment remained comparatively low at Day 1 and showed only modest change by Day 5, indicating limited enrichment of reparative fibroblasts. Similarly, Ki67⁺ cell density was maintained at similar levels from Day 1 to Day 5 and was significantly lower than in +FLS synovial gels at Day 1. Thus, the observed synovial expansion, as indicated by increased total synovial cell density, is primarily driven by migration from the tendon hydrogel **(Supplementary Fig. S3)**.

To determine whether synovial activation influences immune cells recruitment, we next examined the transendothelial migration of CellTracker-labeled monocytes into the synovial and tendon hydrogels using live confocal imaging at Days 1, 3, and 5 (**Fig. 4d**), which demonstrated that the inclusion of FLS in the synovial hydrogel increased monocytes transmigration from the vascular compartment to the tendon compartment. Quantitative analysis of the confocal images confirmed a significant increase in monocyte infiltration for +FLS devices at Days 3 and 5 compared to -FLS devices and revealed a time-dependent increase in infiltrating monocytes, which was not observed in -FLS devices.

### Synovial FLS induce adhesion-like peritendinous matrix deposition independent of TGF-β1

With the observation that FLS drive synovial hyperplasia and inflammation, we next sought to examine the extracellular matrix deposition at the synovial-tendon interface. Macroscopic images of contracted tendon hydrogels revealed remarkable matrix attachments between the synovial hydrogel at the bottom of the vascular barrier and the tendon construct, which in conditions lacking exogenous TGF-β1 appeared to occur almost exclusively in +FLS devices. To better characterize this matrix composition, we acquired confocal images of devices immunostained for collagen III and fibronectin at Day 5.

In the absence of exogenous TGF-β1, we observed fibroblast migration into the initially acellular synovial gel in -FLS devices (**Fig. 5a**). While cells at the edge of the synovial gel expressed fibronectin, their matrix rarely connected with the boundary of the tendon construct (**Fig. 5b**). There was also minimal collagen III expressed by cells in the synovial or tendon gels. In contrast, +FLS devices showed robust collagen III and fibronectin-rich networks spanning the fibrotic tendon and synovial gels, recapitulating adhesion-like bridging networks connecting the two compartments (**Fig. 5c,d**).

**Figure 5.**
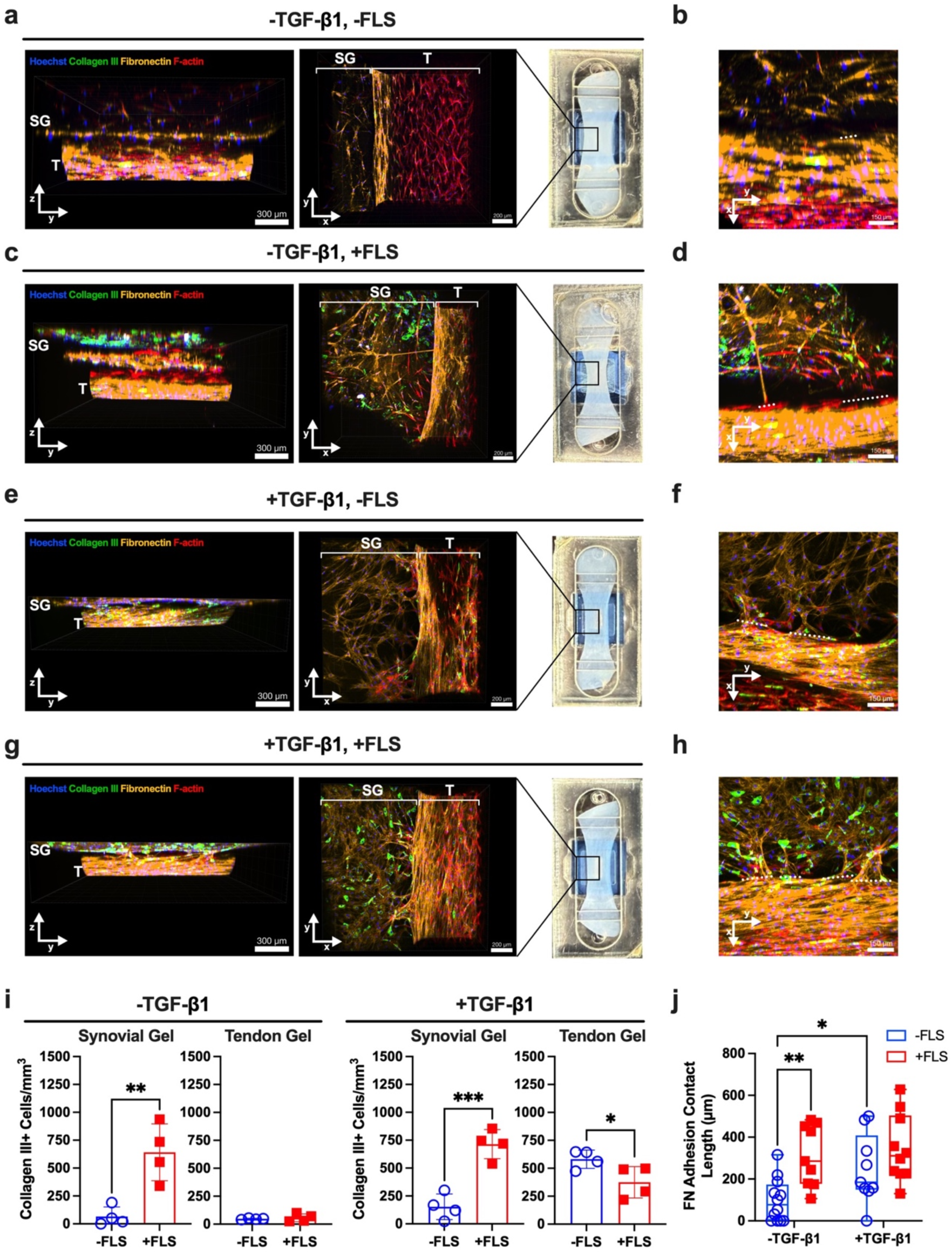
Synovial FLS induce fibronectin- and collagen III-rich matrix bridging and adhesion-like networks at the synovial-tendon interface. **a-d**, Confocal reconstructions of synToC devices cultured without TGF-β1 showing side (left) and top (right) views (SG = synovial gel, T = tendon gel). Devices were stained for collagen III (green) and fibronectin (yellow), with Hoechst (blue) and F-actin (red) as counterstains. The macroscopic device image specifies the location of the confocal image field of view at the synovial-tendon interface. **a,b,** In -FLS devices, fibroblast migration into the synovial gel is observed, but fibronectin networks remain largely compartmentalized with minimal matrix continuity between the synovial and tendon gels. **c,d,** In +FLS devices, dense fibronectin- and collagen III-rich networks bridge the synovial and tendon compartments, forming direct matrix extensions characteristic of adhesion-like contacts. Insets highlight synovial-tendon interface regions with a white dashed line. **e-h,** Confocal images of +TGF-β1 devices at Day 5. Fibronectin-rich interfacial contacts develop in both -FLS and +FLS conditions; however, collagen III deposition within the synovial gel is markedly enriched in +FLS constructs. **i,** Quantification of collagen III immunofluorescence expression in the synovial and tendon gels, reported as the number of collagen III+ cells mm^-3^. Data are mean ± s.d. (n = 4 devices per condition); unpaired t-test; *p<0.05, **p<0.01, ***p<0.001. **j,** Quantification of adhesion contact length, measured as the length of fibronectin signal that contacts both the synovial and tendon gels, parallel to the tendon long axis. Data are mean ± s.d. (n = 4 devices per condition); unpaired t-test; two-way ANOVA with Tukey’s post-hoc test; *p<0.05, ****p<0.0001.

Upon TGF-β1 stimulation, fibronectin-rich ECM networks connecting the synovial and tendon gels formed in both ±FLS devices (**Fig. 5e-h**), indicating that profibrotic signaling can independently induce this adhesion-like phenomenon. However, the density of these fibronectin-rich bridging networks was more pronounced and collagen III expression was more abundant at the synovial-tendon interface of +FLS devices (**Fig. 5g,h**). This observation is consistent with reports that FLS are key contributors to collagen deposition in fibrotic pathologies^16^. Quantification demonstrated that FLS increased collagen III expression in the synovial gel regardless of TGF-β1 stimulation (**Fig. 5i**). In contrast, collagen III expression was low in the tendon gels of -TGF-β1 devices regardless of FLS addition, but was significantly increased with TGF-β1 stimulation (**Fig. 5i**). To quantify adhesion formation, we determined adhesion contact length by measuring the total distance of fibronectin contact between the synovial and tendon gels, consistent with established histological adhesion metrics^41^. Incorporation of FLS led to significantly greater adhesion contact length compared to -FLS devices in the absence of TGF-β1 (**Fig. 5j**). Adhesion contact length was not enhanced further upon exogenous TGF-β1 stimulation.

### FLS and TGF-β1 amplify pro-inflammatory cytokine secretion

To characterize secreted factors in the synovial tendon-on-a-chip under ±FLS and ±TGF-β1 conditions, we analyzed device supernatants at Day 1 and Day 5 using multiplex cytokine profiling. Heatmaps display relative levels of pro-inflammatory cytokines, and reveal that IL-6 and IL-8 were the most abundant cytokines at both timepoints (**Fig. 6**). Hierarchical clustering also indicated that IL-6 and IL-8 have similar response patterns (**Fig. 6a**), suggesting coordinated regulation in adhesion-relevant synovial hyperplasia, proliferation, and leukocyte migration^42, 43^. At Day 1, GM-CSF, TGF-β1, and TGF-β2 were the next most prevalent factors. At Day 5, GM-CSF and TGF-β2 demonstrated a similar response pattern and exhibited overall increases compared to Day 1, while cytokines linked to T cell differentiation and adaptive immune activation, including IL-2, IL-4, IL-5, IL-12p70, and IFN-γ,^44–46^ remained low, consistent with the absence of lymphocytes in this model.

**Figure 6.**
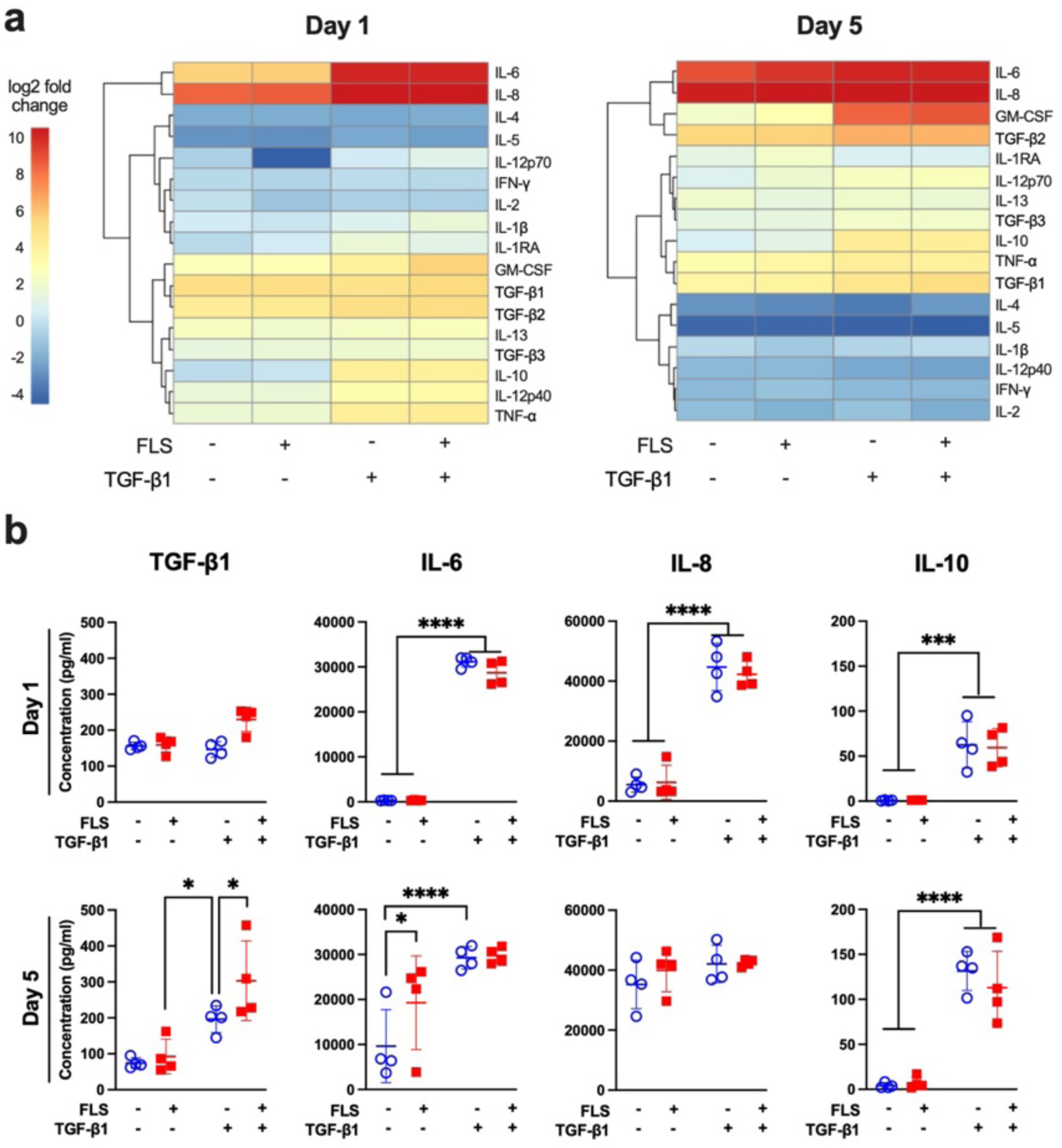
Presence of FLS and TGF-β1 signaling modulate cytokine secretion profiles. **a**, Heatmaps showing cytokine profiles at Days 1 and 5 comparing ±FLS devices cultured with or without TGF-β1. Results are presented as log2 fold change from the -TGF-β1, -FLS baseline condition. Rows are clustered by similar protein expression patterns. At both time points, IL-6 and IL-8 are significantly elevated. **b,** Plots detailing the concentrations of secreted active TGF-β1, IL-6, IL-8, and IL-10 from device supernatants. Data are mean ± s.d. (n = 4 devices per condition); Two-way ANOVA with Tukey’s post-hoc test; *p<0.05, ****p<0.0001.

We next plotted the concentrations of each cytokine to interpret FLS- and TGF-β1-dependent differences between conditions (**Fig. 6b and Supplementary Fig. S4)**. While secreted active TGF-β1 levels were similar among all four conditions at Day 1, we observed significantly higher TGF-β1 levels in +FLS devices at Day 5 (303.1±110.7 pg ml^-1^) compared to -FLS devices (196.1±37.3 pg ml^-1^). Interestingly, this relationship was only true in TGF-β1-stimulated conditions. Our measured active TGF-β1 concentrations are consistent with values reported from human samples in the initial and early stages of peritendinous adhesion formation^1^. At Day 1, exogenous TGF-β1 stimulation led to significant elevation of IL-6, IL-8, and IL-10 for both ±FLS devices. In -TGF-β1 conditions, there was an accumulation of IL-6 (-FLS: 30-fold, +FLS: 100-fold) and IL-8 (±FLS: 6-fold) from Day 1 to Day 5. Conversely, in +TGF-β1 conditions, IL-6 and IL-8 levels were sustained at high concentrations from Day 1 to Day 5, indicating that strong profibrotic signaling drives a maximal inflammatory state independent of time. IL-10 levels were significantly elevated upon TGF-β1 stimulation at both timepoints. At Day 5 in the absence of TGF-β1, FLS drove a significant increase in IL-6 compared to -FLS devices. Given the role of IL-6 in tendon fibroblast activation^47^ and leukocyte recruitment^48, 49^, IL-6 presents a potential target to attenuate FLS-mediated adhesion formation.

### Pharmacological IL-6/JAK/STAT inhibition suppresses inflammatory synovial activation and adhesion formation

Given the central role of IL-6 in tendon fibroblast activation^47^ and leukocyte recruitment^48, 49^, we next tested whether pharmacologic inhibition of the IL-6/JAK/STAT axis could mitigate adhesion-like phenotypes under basal (-TGF-β1) conditions. FLS-laden synToC devices were treated with two FDA-approved drugs targeting the IL-6/JAK/STAT signaling axis. Tocilizumab (TCZ) is a recombinant monoclonal antibody directed against soluble and membrane-bound interleukin 6 receptors (IL-6R)^50^, and tofacitinib (TOFA) is a small molecule inhibitor of JAK1, JAK2, and JAK3^51, 52^ (**Fig. 7a**). We first determined optimal drug doses of 10 µg ml^-1^ for TCZ and 100 ng ml^-1^ for TOFA, which were effective in attenuating nuclear pSTAT3 in FLS monocultures without adversely affecting EC and FLS viability **(Supplementary Fig. S5)**. pSTAT3 is the activated form of the STAT3 transcription factor which acts as a critical regulator of cell growth, survival, differentiation, and inflammation^53^. These concentrations are consistent with median steady-state serum concentrations observed in patients^54, 55^.

**Figure 7.**
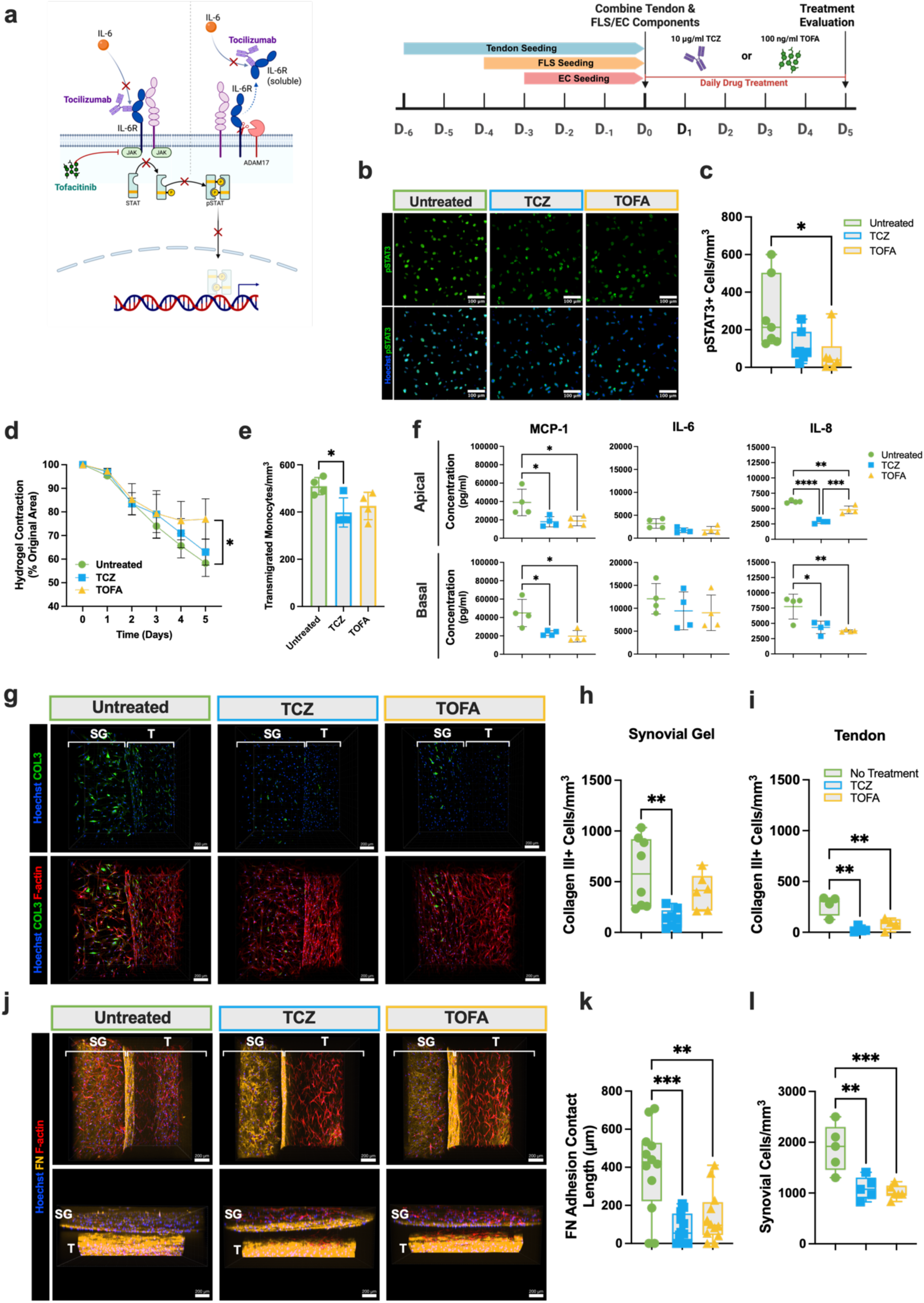
Tocilizumab and tofacitinib attenuate synovial JAK/STAT activation, collagen III deposition, and fibronectin-mediated synovial-tendon adhesions. a, Drug treatment strategy and experimental timeline for the synToC drug testing scheme, outlining the mechanisms and timing of tocilizumab (TCZ) and tofacitinib (TOFA) dosing. b, Nuclear pSTAT3 expression in the synovial gel, with c, quantification demonstrating a significant reduction in pSTAT3 immunofluorescence with TOFA treatment. d, Quantification and representative images of tendon hydrogel contraction from Days 0-5, plotted as the percent of original hydrogel area. t-test at each time point: n=4 devices, *p<0.05. e, Monocyte transmigration at Day 2, reported as the number of monocytes transmigrated per unit volume. f, Secreted cytokine levels measured from Day 5 supernatants sampled from the apical (vascular) and basal (tendon) compartments. One-way ANOVA with Tukey’s post-hoc test: n=4 devices per condition, *p<0.05, **p<0.01, ***p<0.001. g, Immunofluorescent confocal images at Day 5 of synToC culture, with collagen III (green), Hoechst (blue), and F-actin (red). Images compare an untreated control group to TCZ and TOFA treatments. Quantification of the number of collagen III+ cells in the h, synovial gel (SG) and i, tendon (T). TCZ significantly attenuated collagen III expression in the synovial gel, while both treatments significantly decreased collagen III expression in the tendon. One-way ANOVA with Tukey’s post-hoc test: n=8 devices per condition, **p<0.01. j, Confocal image reconstructions of Day 5 synToC devices, showing a top-down view (top row) and a side view (bottom row). Devices were stained for fibronectin (yellow), Hoechst (blue), and F-actin (red) to compare fibronectin adhesion contacts between untreated and treated devices at Day 5. k, Quantification of adhesion contact length. Both TCZ and TOFA treatment significantly reduced the formation of fibronectin contacts between the synovial and tendon gels. One-way ANOVA with Tukey’s post-hoc test: n=12 devices per condition, **p<0.01, ***p<0.001. l, Total cell density in the synovial gel, reported as the number of cells/mm^3^. Untreated synToC devices had a significantly greater number of cells in the synovial gel compared to drug-treated devices. One-way ANOVA with Tukey’s post-hoc test: n=5 devices per condition, **p<0.01, ***p<0.001.

FLS-laden devices were treated with either 10 µg ml^-1^ TCZ or 100 ng ml^-1^ TOFA starting at Day 0, following full assembly of the vascular-synovial and tendon compartments (**Fig. 7a**). Drug treatment media was replaced in the vascular compartment daily from Day 0 to Day 5. Immunofluorescence staining demonstrated reduced pSTAT3 expression within the synovial gel following TCZ or TOFA treatment (**Fig. 7b,c**), although pJAK2 expression was not affected **(Supplementary Fig. S6)**.

TOFA, but not TCZ, significantly reduced tendon hydrogel contraction over time, demonstrating attenuation of intrinsic matrix-driven tissue remodeling (**Fig. 7d**). Analysis of soluble mediators at Day 5 demonstrated a 2-fold reduction in MCP-1 in both apical and basal compartments with TCZ and TOFA treatment (**Fig. 7f**), along with significant suppression of IL-8 in both compartments. IL-6 concentrations remained detectable and were not significantly altered, consistent with receptor blockade rather than cytokine depletion. Consistent with reduced chemokine concentrations, monocyte infiltration was significantly attenuated at Day 2 with TCZ treatment (**Fig. 7e**). At the cellular level, TCZ significantly reduced collagen III deposition within the synovial gel (**Fig. 7g,h**), and both TCZ and TOFA suppressed collagen III accumulation within the tendon compartment (**Fig. 7g,i**). Although fibronectin expression within the synovial gel appeared elevated in the TCZ treated synovial gel (**Fig. 7j**), both TCZ and TOFA significantly reduced the formation of adhesion contacts between the synovial and tendon gels (**Fig. 7k**). Synovial cell density was also reduced in TCZ and TOFA-treated chips, indicating that both treatments attenuated synovial hyperplasia (**Fig. 7l**).

## Discussion

Peritendinous adhesions remain a significant clinical challenge, with no approved therapies that prevent or reverse adhesion formation^6, 56^. Despite decades of investigation using small and large animal models, the cellular and molecular mechanisms that drive adhesion formation remain incompletely understood, necessitating the need for new approach methodologies (NAMs). In this study, we report the first demonstration of peritendinous adhesion formation in a synovial tendon-on-a-chip (synToC). Our findings support a central role for synovial fibroblasts (FLS) in driving adhesion pathology by enhancing tissue contraction, matrix deposition, inflammation driven by abundant IL-6 and IL-8 secretion, and increased monocyte transendothelial tissue infiltration. Importantly, FLS drove the formation of fibronectin- and collagen III-rich networks that connected the tendon and synovial hydrogels, resembling nascent peritendinous adhesions observed clinically and providing a mechanistically informative platform for therapeutic testing. These phenotypes emerged even in the absence of exogenous TGF-β1, indicating the significant role FLS play in the multicellular crosstalk in the synToC. Because clinical range-of-motion outcomes correlate with imaging-detected peritendinous adhesions^25–27^, the synToC, which recapitulates adhesion-like tissue formation *in vitro*, provides a clinically anchored metric for mechanistic studies and preclinical evaluation of anti-adhesion therapies. Within this context, we evaluated the FDA-approved drugs tocilizumab and tofacitinib and found that pharmacologic inhibition of IL-6/JAK/STAT signaling suppressed synovial activation, inflammatory cytokine secretion, matrix deposition, and interfacial adhesion formation. These results demonstrate the utility of the synToC as a human-relevant NAM for mechanism-informed therapeutic testing that may help de-risk translation of anti-adhesion drug candidates in preclinical studies and subsequent clinical trials.

We first established a vascular-synovial interface to model the native tendon sheath microenvironment^28, 57^. Consistent with prior vascular-synovial chip models^58^, FLS retained characteristic synovial phenotypes, including cadherin-11 and PRG4 expression, while endothelial cells formed mature VE-cadherin junctions. By seeding FLS within a collagen gel rather than as a monolayer, mimicking the 3D architecture of synovial ECM, the co-culture exhibited robust FLS proliferation and increased collagen III deposition. This response is in line with *in vivo* observations of post-injury scar tissue and synovial fibrosis^17, 59^. Although TGF-β1 enhanced collagen III expression, FLS proliferation was independent of TGF-β1, indicating a constitutively activated phenotype that may contribute to pathological responses.

Synovial fibroblasts were key regulators of tendon matrix remodeling in the synToC, likely through paracrine signaling within the synovial-tendon microenvironment. In devices stimulated with TGF-β1, tendon contraction was comparable with or without FLS, suggesting that the growth factor is a dominant driver of tissue fibrosis, as we and others have previously shown^1, 19, 60, 61^. In contrast, under basal conditions in the absence of TGF-β1, FLS was sufficient to drive the tendon fibrotic phenotype relative to acellular synovial controls. Measurement of secreted active TGF-β1 in device supernatants revealed elevated levels in FLS-containing devices when stimulated with TGF-β1, implying autocrine and paracrine signaling feedback loops, which might also involve IL-6 signaling. It is also possible that FLS-driven contraction of the tendon hydrogel, potentially through integrin-mediated mechanobiological pathways, enhances sequestration and local activation of TGF-β1 within the matrix^62^. This may reduce detectable soluble levels in the supernatant while promoting a contractile phenotype. However, tissue levels and activity of TGF-β1 were not directly measured in this study. Furthermore, synovial fibroblasts have been shown to amplify fibrotic signaling through paracrine cytokine production and matrix remodeling through α-SMA activation in synovitis, corroborating our findings in the synToC^63^. Interestingly, previous studies have reported that sustained matrix contraction and scar maturation can occur even as α-SMA expression declines^64^, indicating that mature fibrotic tissues may maintain contractile behavior through α-SMA-independent mechanisms, as observed in the synToC at later time points. Our observations are therefore consistent with the concept that synovial fibroblasts modulate tendon fibrosis through both early TGF-β-mediated myofibroblast activation and subsequent matrix remodeling processes, which remain to be elucidated.

Beyond tendon matrix fibrotic remodeling, we demonstrated that synovial fibroblast crosstalk was sufficient to promote deposition of adhesion-like matrix at the tendon-synovial interface. Peritendinous adhesions after tendon injury have been shown to involve both intrinsic fibroblast activity within the tendon and extrinsic cell infiltration from surrounding tissues, including synovium and circulating cells^65^. Further, fibroblast adhesion and chemotaxis shortly after injury have been linked directly to fibronectin and type III collagen secretion^11, 66^. Correspondingly, we observed that adhesion formation in the synToC was associated with fibroblast proliferation, migration, and matrix deposition, even in the absence of exogenous TGF-β1. The expression of CD90, a surface marker associated with fibroblast activation in fibrosis^67, 68^, was predominantly increased and localized to the synovial compartment containing FLS. While tendon-resident fibroblasts migrated externally and deposited fibronectin in the control synToC lacking FLS, CD90 expression and adhesions were less pronounced, consistent with previous reports of reduced CD90 expression in inflammatory-stimulated tendon-resident cells^69^. Consistent with this, fibroblast proliferative activity, indicated by Ki67 expression, was confined to the synovial compartment in +FLS devices, with minimal proliferation among tendon fibroblasts. These findings underscore the ability of the synToC to resolve cell type-specific contributions^65^ to adhesion formation in an in vitro human-relevant NAM, where mechanistic studies have previously relied on animal models.

Adhesion formation in our model was also accompanied by elevated inflammatory cytokines and enhanced monocyte transmigration, corroborating observations that fibrosis is tightly coupled to immune cell recruitment^70, 71^. These observations support the paradigm that stromal cells, such as synovial fibroblasts, play non-canonical roles as immune sentinel cells that produce chemokines and factors that guide immune cell recruitment and activation^72^. To our knowledge, our results provide the first evidence that synovial fibroblasts enhance monocyte transmigration in an in vitro model of intrasynovial tendon microenvironment, making the synToC a suitable model of stromal immunity in synovial diseases. These findings reflect the key role of FLS-derived secreted factors in driving immune cell recruitment and activation. Among these signals, IL-6 emerged as a plausible mediator of adhesions independent of TGF-β1. IL-6, which is upregulated in peritendinous adhesion tissue^73–75^, promotes cell proliferation, matrix secretion, and angiogenesis via JAK/STAT signaling^76, 77^. Given that CD90+ fibroblasts have been identified as primary sources of IL-6^78^, our results support the hypothesis of a feed-forward loop in which synovial fibroblasts amplify IL-6 signaling to promote adhesion formation, and identify IL-6 as a therapeutic target for tendon adhesions.

Although TGF-β1 is a well-established driver of adhesions^1^, its pleiotropic role in tendon repair rules out its inhibition as a therapeutic option^60, 61^. On the other hand, the IL-6/JAK/STAT pathway represents a clinically actionable target for tendon and post-operative adhesions, as has been demonstrated in animal studies using experimental inhibitors of IL-6^79–81^. Remarkably, tocilizumab and tofacitinib attenuated key adhesion-associated readouts in the synToC, including collagen III deposition in both the synovial and tendon gels, synovial hyperplasia, inflammatory cytokine secretion, and most importantly fibronectin-rich adhesion contacts. As expected, IL-6 levels remained unchanged^82^, while both treatments reduced MCP-1 and IL-8, implicating diminished immune cell recruitment as an underlying anti-adhesion mechanism. The synToC corroborates therapeutic interventions in animal studies and provides human-relevant evidence that positions JAK/STAT blockade as a viable clinical target to prevent peritendinous adhesions following flexor tendon repair or tenolysis surgery.

There are several limitations to this study, including the use of primary cells from different sources, which introduces biological variability into the model. Nevertheless, the fibrotic adhesion phenotypes were evident and reproducible against this donor variability. In particular, the tendon cells we used, which were isolated from fibrotic peritendinous surgical waste tissue, may retain epigenetic and transcriptional memory of prior inflammation, potentially biasing their behavior toward a pro-fibrotic state. While this makes them unsuitable for modeling of healthy tendon biology, they are still appropriate for studying the pathological microenvironment associated with peritendinous adhesions. Furthermore, the use of joint-derived rather than tendon sheath-derived synovial fibroblasts is another limitation. However, previous studies indicate broadly similar proliferative and inflammatory behavior across synovial fibroblast sources^83, 84^. The simple approach used in cell sourcing for the synToC may not necessarily capture the cellular complexity in adhesions. Future studies could incorporate single-cell transcriptomic profiling to resolve fibroblast, immune, and endothelial cell heterogeneity in the synToC and map these populations onto emerging cellular atlases of human tendon adhesion tissue^40^.

In conclusion, we establish a synovial tendon-on-a-chip (synToC) as a new approach methodology that bridges the gap between reductionist in vitro models or animal injury models and testing in humans. Importantly, the synToC recapitulates key features of the intrasynovial tendon microenvironment including synovial hyperplasia, immune cell recruitment, and adhesion-like tissue formation in vitro. The synToC provides quantitative functional readouts in a controllable microphysiological system, enabling mechanistic study of multicellular crosstalk in adhesion pathogenesis and preclinical evaluation of anti-adhesion therapies.

## Methods

### Cell culture details

Primary human fibroblast-like synoviocytes (FLS) were obtained from Cell Applications (San Diego, CA). Tissue donor was a 63-year-old Caucasian male and cells were isolated from normal synovial tissue. FLS were expanded in T-75 flasks in Synoviocyte Growth Medium (Cell Applications, 415K-500). Tendon fibroblasts were isolated from tenolysis tissue samples under an approved University of Rochester Institutional Review Board protocol (STUDY00004840). Tissue donor was an 80-year old Caucasian male and cells were isolated from the left ring digital flexor tendon. Tendon fibroblasts were expanded in T-175 flasks in MEM Alpha media (Gibco, 12571063) supplemented with 10% FBS and 1% Penicillin-Streptomycin and verified for scleraxis expression using immunofluorescence. Cryopreserved pooled human umbilical vein endothelial cells (HUVECs) were purchased from Vec Technologies Inc. and expanded in T-25 flasks (Corning Inc., 430,639). HUVECs were cultured in EGM-2 Endothelial Cell Growth Medium (Lonza, CC-3162). All cells were maintained under standard culture conditions (37°C, 5% CO_2_, 95% humidity) and used between passages 2 and 6.

### Peripheral blood mononuclear cell (PBMC) isolation and CD14 + monocyte sorting

PBMCs were isolated fresh before each experiment from healthy donors with informed consent according to an approved protocol by the University of Rochester Institutional Review Board (STUDY0004777). All methods were performed in accordance with relevant guidelines and regulations. First, PBMCs were isolated from whole blood samples using SepMate™ according to the manufacturer instructions (Stem Cell Technologies, 85460). The enriched PBMCs were washed with 1X DPBS + 2% FBS twice by centrifuging at 300 x g for 8 minutes at room temperature. After removing the supernatant, the cell pellet was resuspended in isolation buffer consisting of 1X DPBS (Gibco, 14190250), 2 mM EDTA (Gibco, 15575020), and 1 mg ml^-1^ BSA. The cell suspension was filtered through a 70 µm cell strainer, spun down at 300 x g for 3 min, and suspended in 80 µl of isolation buffer. To sort for CD14+ monocytes, CD14 microbeads (Milltenyi Biotec, 130-050-201) were added to the cell suspension and incubated at 4°C for 15 min. The cell suspension was again centrifuged at 300 x g for 3 min and the cells were suspended in 500 µl of isolation buffer. A QuadroMACS™ separator was assembled with an LS column (Milltenyi Biotec, 130-042-401) and the 500 µl cell suspension was added to the column. The column was washed 3 times with 3 ml of isolation per wash. CD14+ monocytes were then flushed out of the column and used in experiments within 3 h of blood collection.

### synToC device components and nanomembranes

The microfluidic synToC device consists of two acrylic components as well as an accessory reservoir component (18 cm length x 9 cm width) manufactured by ALine Inc. (Signal Hill, CA) separated by a 3-µm dual-scale nanomembrane. Silicon nitride nanomembranes manufactured by SiMPore, Inc. (Henrietta, NY) and serve as biocompatible substrates for cell culture. 3 µm dual-scale membranes (∼100 nm thick, ∼60 nm diameter nanopores, 3 µm diameter micropores, ∼15% nanoporosity, ∼0.7% microporosity) were used in this study. We have previously validated that 3 µm dual-scale membranes permit monocyte transmigration^24^. Components were assembled in sterile conditions in a biological safety cabinet, as previously described^19, 85^. Briefly, a membrane chip was placed on a custom assembly jig in a non-inverted orientation using notched chip tweezers. This results in a trench-up chip configuration for the final assembled device. The protective layers surrounding the top component were removed to expose a pressure-sensitive adhesive (PSA). The top component was placed over the nanomembrane and pressed firmly with the jig to activate the PSA. The outer protective layers were removed from the bottom component, while keeping the final masking layer intact to keep the PSA covered. The bottom component and reservoir were placed in a petri dish alongside the assembled top component and membrane. All components were sterilized under UV for 20 minutes.

### synToC culture timeline

The synovial integrated was prepared as outlined in Figure 1a. First, tendon fibroblasts were lifted from flasks by rinsing with 1X DPBS and applying 0.25% trypsin-EDTA for 3 min. Tendon fibroblasts and freshly isolated monocytes were encapsulated in a type I collagen hydrogel (Advanced Biomatrix, 5026) at a 7 to 1 ratio of tendon fibroblasts to monocytes (tendon fibroblast cell density: 400,000 cells ml^-1^). 100 µl of the hydrogel cell suspension was added to the bottom component and was allowed to crosslink in the incubator for 20 min. The protective layer was then removed from the reservoir component, and the reservoir was attached to the bottom component. The hydrogels were scored on the long edges between anchors to enable contraction throughout the culture period. 200 µl of X-VIVO 10 media (Lonza, 04-380Q) supplemented with 20 ng ml^-1^ M-CSF (PeproTech, 300-25) was added to each reservoir. Hydrogels were cultured for 6 days (D_-6_-D_0_), with media half-replacements occurring each day. At D_-4_, top components were prepared by rinsing the wells with 100 µl of 1X DPBS. The top side of the nanomebrane was coated with fibronectin at 5 µg/cm^2^ for 1 h at room temperature. The coating solution was removed and replaced with 50 µl of Synoviocyte Growth Medium. FLS were lifted from flasks by first rinsing with HBSS (Gibco, 14175095) and applying 0.25% trypsin-EDTA for 3 min. FLS were encapsulated in a type I collagen hydrogel (Advanced Biomatrix, 5026) at a density of 100,000 cells ml^-1^. 15 µl of the FLS gel was added to the backside of the nanomembrane. In the case of -FLS devices, 15 µl of hydrogel without FLS was added instead. The devices were kept in an inverted position while the hydrogels crosslinked in the incubator for 15 min. An additional petri dish was prepared by adhering reservoir components on to the surface of the dish and filling the reservoirs with 200 µl of Synoviocyte Growth Medium. The top components with crosslinked hydrogels were flipped onto the media reservoirs, with the open well facing up. After 24 h of culture, HUVECs were lifted from flasks by rinsing with 1X DPBS and applying 0.25% trypsin-EDTA for 3 min. Media in the reservoirs was replaced with a 1:1 ratio of Synoviocyte Growth Medium:EGM-2 (FLS-EC co-culture media). HUVECs were seeded on top of the membrane at a density of 130,000 cells ml^-1^. At D_-2_ and D_-1_, media was replaced in reservoirs and top wells with a 1:1 ratio of X-VIVO 10 media:FLS-EC co-culture media. At D_0_, the top vascular-synovial component and the bottom tendon component were combined by first removing the reservoir from the bottom component. Monocytes were freshly isolated and added to the top well at a concentration of 100,000 cells ml^-1^ (10,000 monocytes/device). X-VIVO 10 media with or without 10 ng ml^-1^ TGF-β1 was added to the bottom channel. Experimental analyses were performed from D_1_-D_5_. While we previously demonstrated the impact of circulating flow on the vascular inflammatory response^24^, devices in this study were maintained in static culture conditions to focus on the synovial response and to benchmark against previous static hToC phenotypes^19^.

### Drug treatments

For therapeutic drug treatment experiments, tocilizumab (Selleck Chemicals, A2012) and tofacitinib citrate (Selleck Chemicals, S5001) were prepared by serial dilution in X-VIVO 10 media. A preliminary efficacy study was performed to determine the doses that resulted in the greatest reduction in nuclear pSTAT3 expression compared to an untreated vehicle control. The vehicle control for tocilizumab was 0.75% DPBS and the vehicle control for tofacitinib was 0.01% DMSO. Tocilizumab concentrations included 0.1, 1, 10, and 100 μg ml^-1^ and tofacitinib concentrations included 0.1, 1, 10, and 100 ng ml^-1^. A combination treatment of 10 μg ml^-1^ tocilizumab + 10 ng ml^-1^ tofacitinib was also tested. FLS were cultured in monolayers in a 96 well plate at a density of 70,000 cells ml^-1^ in serum-free X-VIVO 10 media. All conditions except the negative vehicle control were incubated with 10 ng ml^-1^ IL-6 (R&D Systems, 206-IL-010) + respective drug treatment for 30 min. Cells were then promptly fixed with 4% paraformaldehyde and labeled with pSTAT3 via immunofluorescence. For synToC drug treatment experiments, 10 μg ml^-1^ tocilizumab or 10 ng ml^-1^ tofacitinib was added to the top vascular channel at D_0_ and half-replacements were performed each day until D_5_ by removing 50 μl of media from the top channel and adding 50 μl of fresh drug treatment.

### Cell viability assays

To compare FLS cell viability in different media conditions, a LIVE/DEAD™ Viability/Cytotoxicity Kit (Invitrogen, L3224) was used according to the manufacturer’s protocol. FLS were encapsulated in a type I collagen hydrogel at a density of 100,000 cells ml^-1^ and cultured in a 96 well-plate, with 30 µl of hydrogel per well. X-VIVO 10, Synoviocyte Growth Medium, FLS-EC co-culture media, or a 1:1 ratio of X-VIVO 10 media:FLS-EC co-culture media was applied, and cell viability was determined after 1, 3, and 6 days of culture. A Cell Counting Kit-8 assay (Tocris Biosciences, 7368) was used to determine cell viability of FLS and HUVECs cultured in their respective medias supplemented with tocilizumab (0.1-100 µg ml^-1^) or tofacitinib (0.1-100 ng ml^-1^) treatment. Percent cell viability was calculated as the experimental absorbance divided by the vehicle control absorbance.

### Live cell tracking and monocyte infiltration experiment

CellTracker Green CMFDA (Thermo Fisher, C2925), CellTracker Orange CMTMR (Thermo Fisher, C2927), and CellTracker Deep Red (Thermo Fisher, C34565) were used to visualize the spatial localization of tendon fibroblasts, FLS, and monocytes, respectively, within the synToC device. To capture monocyte infiltration from the vascular side to the tendon compartment, monocytes were labeled with CellTracker Deep Red and imaged at Days 1, 3, and 5. Confocal z-stack images were acquired with an Andor spinning disk confocal microscope (Andor Technology, United Kingdom) on Fusion software with a 10x objective and a 4 µm step size. Devies were placed on an incubation stage at standard conditions for the entire imaging session to prevent cell death during imaging. Images were processed in Imaris (Oxford Instruments, United Kingdom).

### Immunofluorescence microscopy

To prepare devices for immunofluorescence staining, devices were first washed with DPBS. The top and bottom channels were fixed with 4% paraformaldehyde (Invitrogen, I28800) for 10 min. Top and bottom channels were washed three times with DPBS, and then blocked for 1h at room temperature with DBPS containing 3% BSA + 0.3% Triton X-100. Following another wash step in DPBS, primary antibodies were diluted in DBPS containing 3% BSA. Primary antibodies were added to the top and bottom channels and incubated overnight at 4°C. Devices were then washed with DPBS 3 times as follows: 200 µl was flushed through bottom channel 2 times, then 50 µl fresh DPBS was added to the top and bottom channel for 20 min. A second blocking step was carried out with 10% goat serum for 30 min at room temperature. Secondary antibodies were diluted in 5% goat serum and incubated in devices for 2 h at room temperature. Devices were then washed with DPBS 4 times in the same manner as the primary antibody washes. Hoechst nuclear stain was added for 5 min, followed by 2 more DPBS washes of the top and bottom channels. Detailed primary and secondary antibody information is included in **Supplementary Table S1**. Images were acquired with an Andor spinning disk confocal microscope on Fusion software with a 10x objective and a 4 µm step size. Images were processed in Imaris and the same brightness and contrast settings were kept consistent across all images of the same marker.

### Second harmonic generation (SHG) microscopy

To visualize collagen fibers within the tendon hydrogels, multiphoton images were acquired using an Olympus FVMPE-RS multi-photon microscope with a 25X (NA 1.5) objective (Olympus Scientific, Waltham, MA). Devices were illuminated with 800-nm light generated by a Mai Tai HP Deep See Ti:Sa laser (Spectra-Physics, Santa Clara, CA) and collagen was visualized using second harmonic generation. Bandpass filters of 370-410 nm (collagen, SHG) or 575-630 nm (AF567-phalloidin) were applied. Images were acquired beginning 50 µm below the surface of the hydrogel with a 1.25x zoom and a 5 µm z-step size.

### Image analysis and quantification

#### Hydrogel contraction quantification

Devices were flipped so the bottom channel was facing up, and a macroscopic image of the tendon hydrogel was captured each day from D_0_-D_5_. Images were imported into Fiji and measurements of the hydrogel area and the rectangular area between the bottom channel posts were taken using built-in tools. The percent hydrogel contraction is reported as the percent of original area and plotted using GraphPad Prism.

#### SHG signal quantification

SHG signal intensity was measured in FIJI from 3 fields of view in 4 devices per condition at the same z-height. Mean intensity per condition was plotted using GraphPad Prism.

#### Percent positive α-SMA and Ki67 cells in tendon hydrogel

Immunofluorescence images of α-SMA and Ki67 in the tenon hydrogel were analyzed in Imaris. The Surfaces object detection tool was used to generate a 3D render of the cell bodies based on F-actin labeling. The F-actin surface served as a mask for intensity classification for proteins of interest. A positive classification threshold was determined for both α-SMA and Ki67 and kept consistent throughout analysis. The percent positive expressing cells was calculated by dividing the number of positive cells by the total number of cells. Mean values were determined per image based on two fields of view in 4 devices and plotted in Graphpad Prism.

#### Positive cell density of collagen III, CD90, Ki67, and pSTAT3

Immunofluorescence images of collagen III, Ki67, and CD90 were analyzed in Imaris. F-actin surfaces were generated for collagen III and CD90 analysis, while Hoechst (nuclear) surfaces were generated for Ki67 and pSTAT3 analysis. A positive classification threshold was determined for each protein and kept consistent throughout analysis. Positive cell density was calculated by dividing the number of positive cells by the total volume of the synovial gel or tendon gel. Mean values were determined per image based on two fields of view in 4 devices and plotted in Graphpad Prism.

#### Synovial cell density

The Spots object detection tool in Imaris was used to identify nuclei based on Hoechst staining intensity. The number of cells in two fields of view in 4 devices was quantified and cell density was calculated by dividing the cell count by the synovial gel volume.

#### Adhesion contact length

A Surface was generated for fibronectin immunofluorescence signal in Imaris. Using built-in measuring tools, the length of the fibronectin surface that contacted both the synovial and tendon gels was measured in 4 devices per condition and plotted in Graphpad Prism.

#### Monocyte infiltration

Confocal images collected on experimental Days 1, 3, and 5 were imported into Imaris for three-dimensional visualization and quantitative analysis of transmigrated monocytes. Images were centered and oriented to ensure uniformity across the images, with the membrane positioned as the upper layer of the stack. Devices were divided into two regions of interest: the membrane and the tissue compartment. Monocyte quantification was performed using the Imaris Spots detection tool. Automatic spot detection was applied, followed by adjustment of the minimum and maximum quality thresholds as needed to accurately identify individual cells. The final quality threshold value used for each image was kept consistent across all images. The total number of transmigrated monocytes mm^-3^ was calculated by dividing the number of transmigrated monocytes by the volume of the tissue compartment. Quantification was recorded for 4 devices per condition and plotted in Graphpad Prism.

### Media sampling & cytokine analysis

Secreted cytokine levels were quantified using a multiplex fluorescent bead assays. To compare cytokine levels in devices ±TGF-β1 and ±FLS, 50 µl of media was sampled from the top vascular channel and diluted 1:2 in X-VIVO 10 media. Media supernatants were stored at −80°C until shipment on dry ice to Eve Technologies (Calgary, Alberta, Canada). Supernatants were analyzed with the Human Cytokine Proinflammatory Focused 15-Plex Discovery Assay® Array (HDF15) and the TGFB 3-Plex Discovery Assay® Multi Species Array (TGFβ1-3). For the synToC drug treatment experiment, media was sampled from the top vascular channel (apical) as detailed above. Additionally, media was sampled from the bottom channel (basal) by adding 200 µl X-VIVO 10 media to a pipette tip attached to one port, acting as a reservoir. 200 µl of media, including the ∼50 µL bottom channel volume, was collected by reverse pipetting from the opposite port. Apical and basal supernatants were analyzed with a Luminex® Discovery Assay (R&D Systems) which included MCP-1, IL-6, and IL-8.

### Statistical Analysis

All experiments were performed with N = 3-4 biological replicates and statistically analyzed with GraphPad Prism software (GraphPad, La Jolla, CA). An unpaired t-test was used to compare differences between two groups and an ordinary one-way ANOVA with Tukey’s post-hoc test was used for comparison of multiple groups, with p < 0.05 considered statistically significant. Grouped data were analyzed via a two-way ANOVA with Tukey’s post-hoc test. All quantifiable data are reported as the mean ± standard deviation.

## Data Availability

All data related to the current study are available from the corresponding author upon reasonable request.

## Acknowledgements

I.L., A.C., A.O., N.G., B.L.M., H.A.A., and J.L.M were supported by NIH UG3TR003281, UH3TR003281, and U2CAG088071. I.L. was also supported by the National Science Foundation Graduate Research Fellowship. Schematics were created with BioRender.com.

## Author Contributions

Conceptualization: I.L., J.L.M., H.A.A. Experiments and Data Acquisition: I.L., A.C., A.O. Data Analysis and Interpretation: I.L., A.C., A.O., N.G., J.L.M., H.A.A. Writing Original Draft: I.L. Writing Review and Editing of Final Draft: I.L., B.L.M, J.L.M, H.A.A. Funding: B.L.M, J.L.M. H.A.A. All authors have read and approved the final submitted manuscript.

## Ethics Approval

Monocytes were obtained from healthy donors with informed consent under an IRB-approved protocol at the University of Rochester (STUDY0004777). Tendon fibroblasts were isolated from tendon tissues obtained during hand surgeries with informed consent under an IRB-approved protocol at the University of Rochester (STUDY00004840). All methods were performed in accordance with relevant guidelines and regulations.

---

**Supplemental Figure S1.**
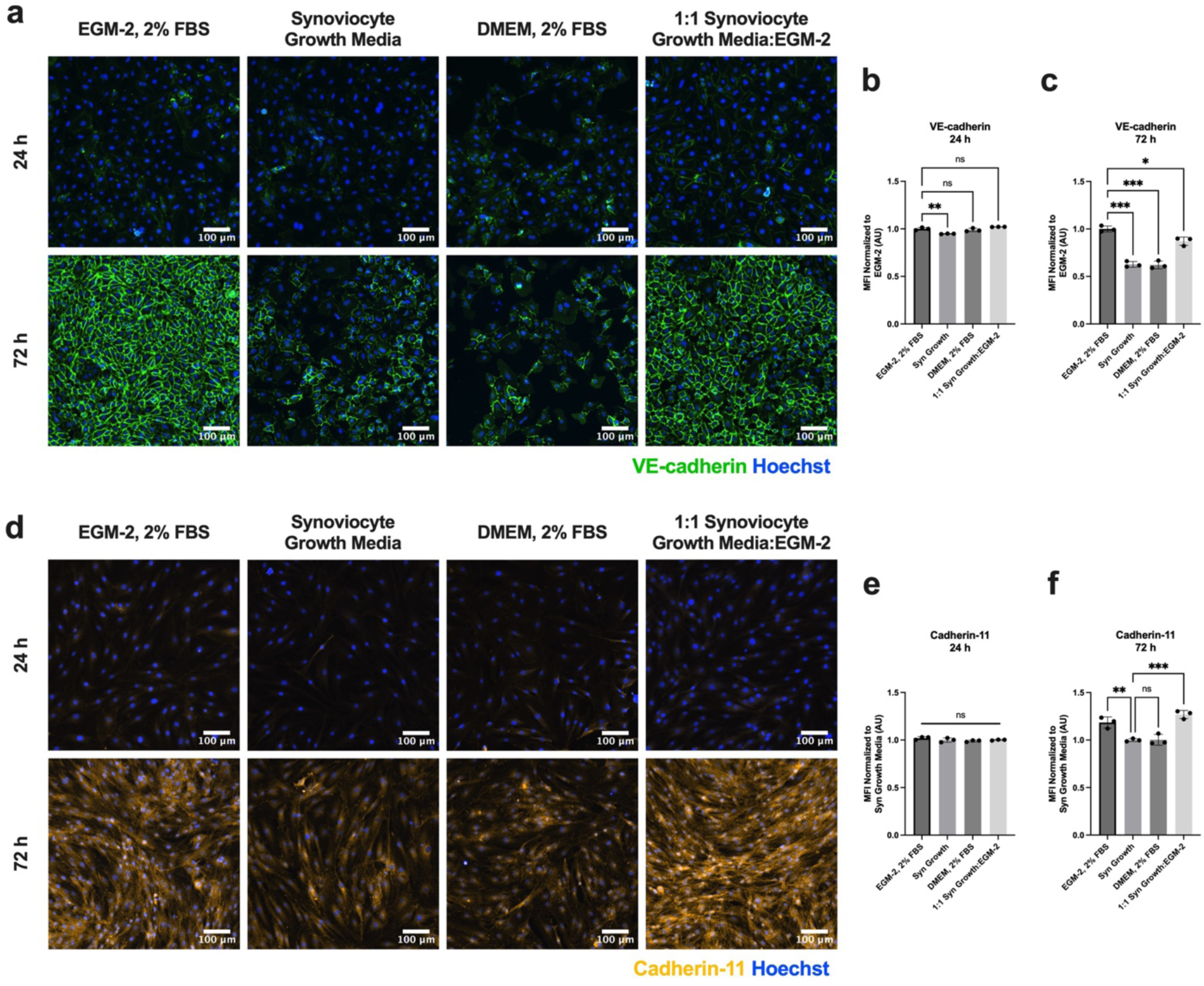
FLS-EC co-culture media optimization - 1:1 ratio of Synoviocyte Growth Media:EGM-2 preserves endothelial VE-cadherin junctions and synoviocyte cadherin-11 expression in 2D culture. **a**, Immunofluorescence images of HUVEC cultured in monolayer and stained for VE-cadherin (green) and Hoechst (blue) compare four different media conditions at 24 h and 72 h of culture. **b,** Mean fluorescence intensity (MFI) of VE-cadherin expression at **b,** 24 h and **c,** 72 h, normalized to MFI of EGM-2 images. Synoviocyte growth media had significantly less VE-cadherin expression at 24 h compared to EGM-2. VE-cadherin MFI for 1:1 Synoviocyte Growth Media:EGM-2 most closely matched that of EGM-2 conditions. **d,** Immunofluorescence images of FLS cultured in a monolayer, stained for cadherin-11 (yellow) and Hoechst (blue) compare the four different media conditions at 24 h and 72 h of culture. **e,** MFI quantification of cadherin-11 expression, normalized to MFI of Synoviocyte Growth Media, demonstrated no significant differences between culture conditions at 24 h. **f,** At 72 h, EGM-2 and 1:1 Synoviocyte Growth Media:EGM-2 had significantly higher cadherin-11 expression compared to Synoviocyte Growth Media, while DMEM had comparable levels. One-way ANOVA with Tukey’s post-hoc test: n=3 wells per condition, *p<0.05, **p<0.01, ***p<0.001.

**Supplemental Figure S2.**
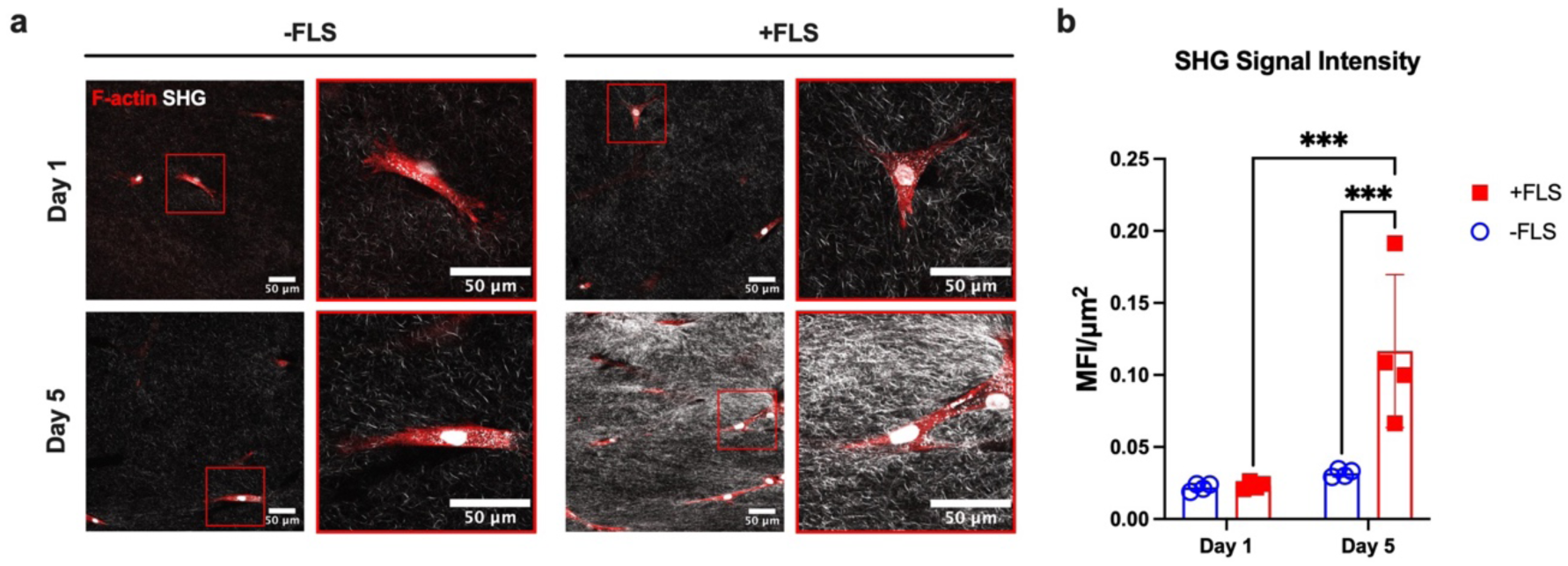
FLS induce significant increases in tendon hydrogel collagen density by Day 5. **a**, Second harmonic generation (SHG) images of tendon hydrogels under basal conditions (-TGF-β1) at D_1_ and D_5_. Tendon fibroblasts are stained with F-actin (red). Red-boxed insets show a zoomed field of view around individual cells. **b,** SHG signal intensity quantification shows significantly heightened SHG signal at D_5_ in +FLS devices. Two-way ANOVA with Tukey’s post-hoc test: n=4 devices per condition, ***p<0.001.

**Supplemental Figure S3.**
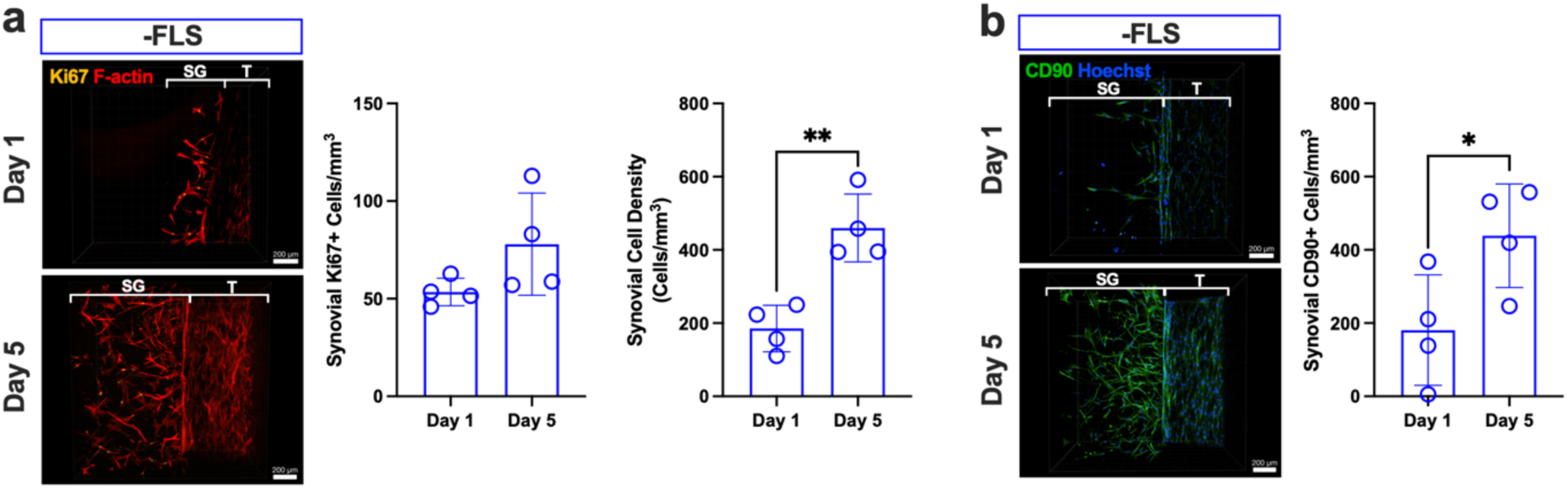
Synovial proliferation and CD90 expression -FLS devices under basal conditions. **a**, Representative confocal images of Ki67 (yellow) and F-actin (red) in the synovial gel (SG) adjacent to the tendon construct (T) at Days 1 and 5 in -FLS devices without exogenous TGF-β1. Quantification of Ki67+ cells within the synovial gel and total synovial cell density, reported as cells mm^-3^. **b,** Representative confocal images of CD90 (green) and Hoechst (blue) in the synovial gel at Days 1 and 5 (-FLS). Quantification of CD90⁺ cells within the synovial gel, demonstrating increased CD90⁺ fibroblast density at Day 5. Data are mean ± s.d. (n = 4 devices per condition); unpaired t-test; *p<0.05, **p<0.01.

**Supplementary Figure S4.**
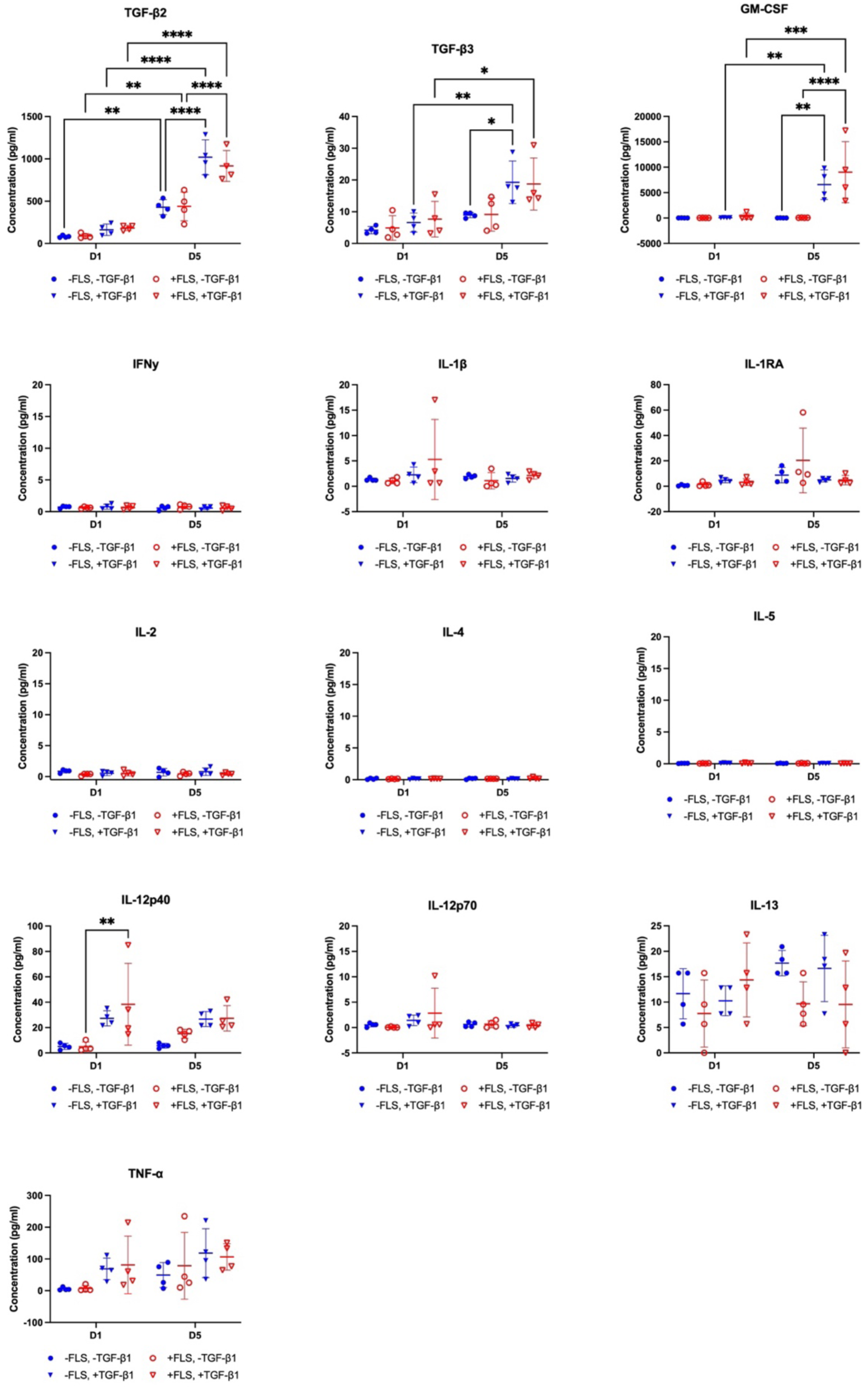
Detailed cytokine expression profiles from sampled supernatants assayed via multiplex analysis. Each plot contains data from devices cultured ±FLS and ±TGF-β1 at Days 1 and 5. All device conditions had significantly elevated TGF-β2 levels from Day 1 to Day 5. TGF-β1-stimulated devices had significantly higher levels of TGF-β2 at Day 5 compared to non-stimulated devices. For +FLS devices, TGF-β3 levels were significantly elevated from Day 1 to Day 5, regardless of TGF-β1 stimulation. GM-CSF levels remained low across all conditions at Day 1 and for -TGF- β1 devices at Day 5. TGF-β1-stimulated devices had significantly elevated GM-CSF levels at Day 5. IFNγ, IL-1β, IL-1RA, IL-2, IL-4, IL-5, and IL-12p70 overall had levels <5 pg ml^-1^ across all conditions. In +FLS devices, IL-12p40 increased with TGF-β1 stimulation at Day 1. IL-13 and TNF-α demonstrated no distinct trends among conditions.

**Supplementary Figure S5.**
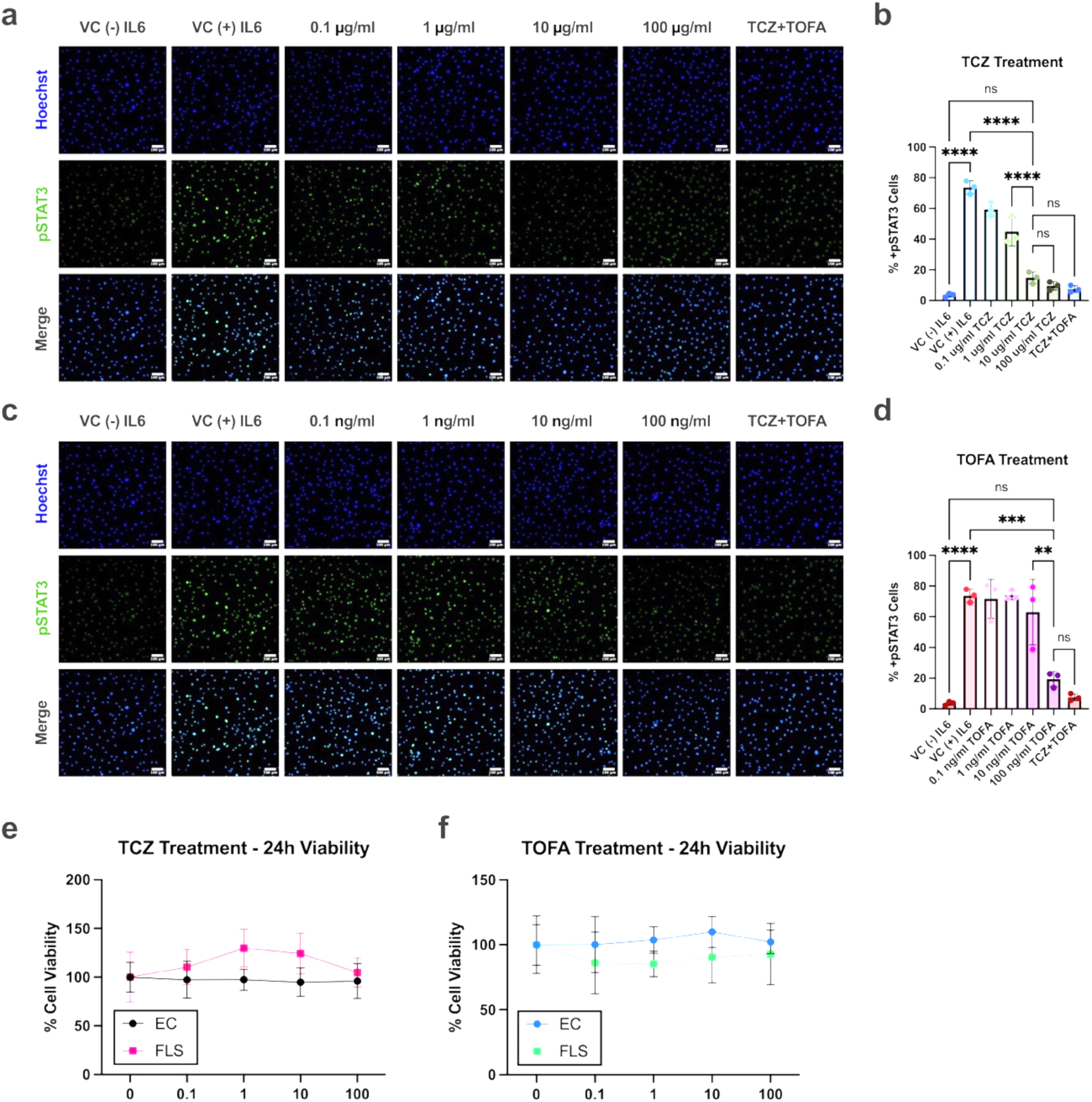
Tocilizumab dose-dependently inhibits IL-6-induced STAT3 activation in FLS while preserving FLS and endothelial viability across clinically relevant concentrations. **a**, Immunofluorescence images of FLS cultured in a monolayer showing nuclear pSTAT3 expression (green), with Hoechst (blue) as nuclear counterstain. FLS were stimulated with IL-6 and treated with a range of tocilizumab (TCZ) concentrations (0.1 μg ml^-1^-100 μg ml^-1^). pSTAT3 expression was compared to a negative vehicle control (VC) without IL-6 stimulation as well as a positive VC with IL-6 stimulation. A 10 μg ml^-1^ TCZ + 10 ng ml^-1^ tofacitinib (TOFA) treatment was also introduced, based on common dose concentrations found in the literature. **b,** Image quantification of panel a, reported as the percent pSTAT3+ cells. Nuclear pSTAT3 demonstrated a dose response to TCZ treatment. The most significant drop in pSTAT3 expression occurred from 1 μg ml^-1^ to 10 μg ml^-1^ TCZ, while there are no significant differences between 10 μg ml^-1^ and 100 μg ml^-1^ or 10 μg ml^-1^ and combination treatment. **c,** Nuclear pSTAT3 expression in FLS stimulated with IL-6 and treated with 0.1 ng ml^-1^-100 ng ml^-1^ TOFA. **d,** TOFA had no impact on pSTAT3 levels until the highest dose of 100 ng ml^-1^. Combination treatment was not significantly different than 100 ng ml^-1^ TOFA. One-way ANOVA with Tukey’s post-hoc test: n=3 wells per condition, **p<0.01, ***p<0.001, ****p<0.0001. **e,** FLS and EC cell viability determined by CCK8 assay after 24 h of TCZ treatments. **f,** FLS and EC cell viability determined by CCK8 assay after 24 h of TOFA treatments. No significant differences in cell viability between untreated controls and TCZ or TOFA doses.

**Supplementary Figure S6.**
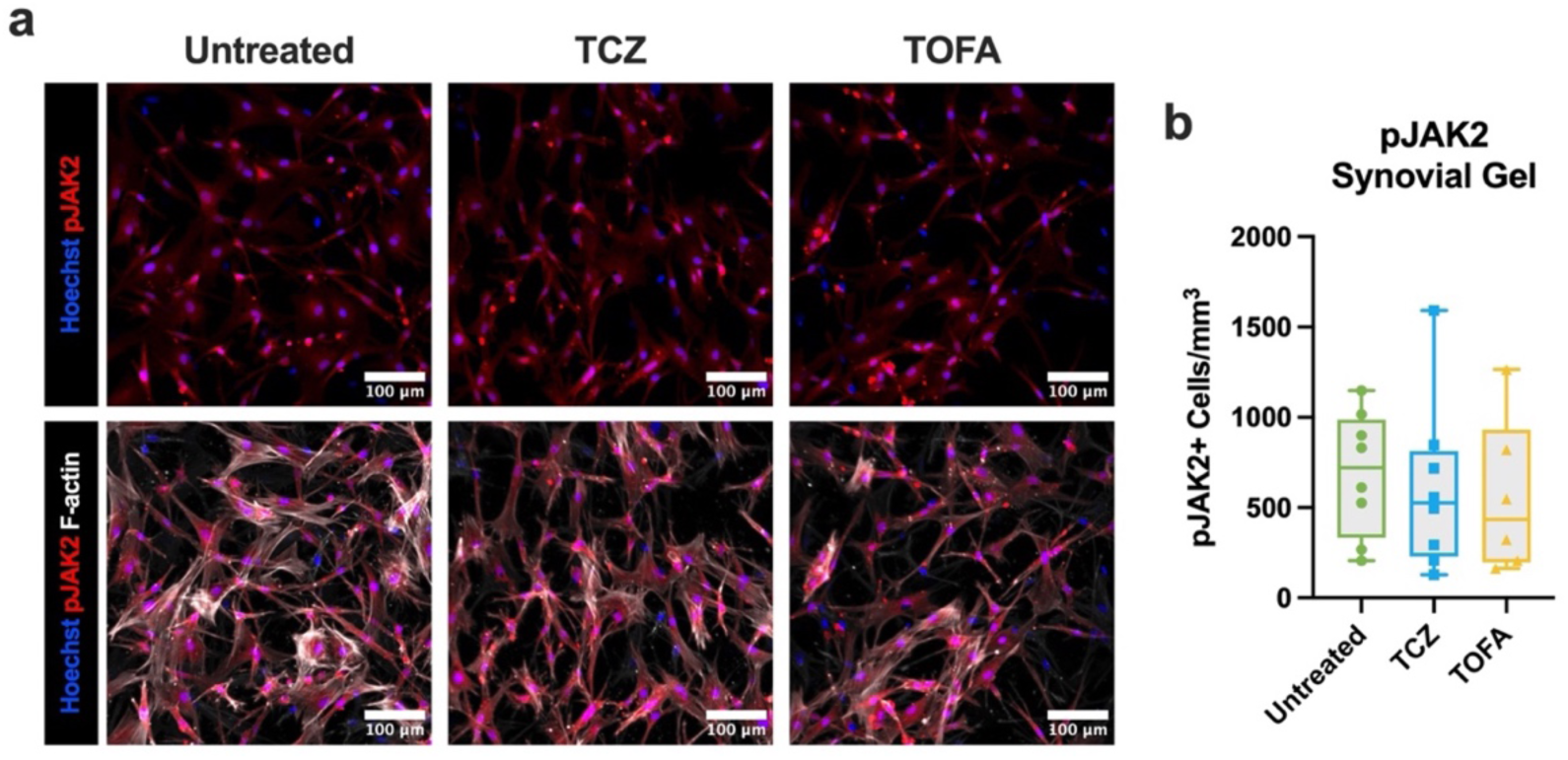
Tocilizumab and tofacitinib treatment do not alter synovial JAK2 phosphorylation in +FLS devices at Day 5. **a**, Day 5 immunofluorescence images of the synovial gels of -TGF- β1, +FLS devices treated with either 10 μg ml^-1^ TCZ or 100 ng ml^-1^ TOFA compared to untreated controls. Devices were stained pJAK2 (red), Hoechst (blue), and F-actin (white). **b,** Quantification of cytoplasmic pJAK2 expression, plotted as the number of pJAK2+ cells mm^-3^. pJAK2 expression was consistent among all treatment conditions.

**Table S1.** Antibodies for immunofluorescence staining.

| Primary Antibody | Dilution | Conjugation | Manufacturer | Catalog # | Secondary Antibody | Dilution | Manufacturer | Catalog # |
| --- | --- | --- | --- | --- | --- | --- | --- | --- |
| **VE-cadherin** | 1:100 | N/A | R&D Systems | MAB8381 | Alexa Fluor 488 | 1:200 | Thermo Fisher | A11001 |
| **Cadherin-11** | 1:100 | N/A | Thermo Fisher | 71-7600 | Alexa Fluor 568 | 1:200 | Thermo Fisher | A11011 |
| **PRG4** | 1:200 | N/A | Sigma | MABT401 | Alexa Fluor 488 | 1:200 | Thermo Fisher | A11001 |
| **Collagen III** | 1:500 | N/A | Abcam | ab6310 | Alexa Fluor 488 | 1:400 | Thermo Fisher | A11001 |
| **Ki67** | 1:1000 | N/A | Abcam | ab15580 | Alexa Fluor 568 | 1:400 | Thermo Fisher | A11011 |
| **PDPN** | 1:100 | N/A | Abcam | ab128994 | Alexa Fluor 568 | 1:400 | Thermo Fisher | A11011 |
| **α-SMA** | 1:1000 | Alexa Fluor 488 | Sigma | F3777 |  |  |  |  |
| **Fibronectin** | 1:500 | N/A | Abcam | ab2413 | Alexa Fluor 568 | 1:400 | Thermo Fisher | A11011 |
| **CD90** | 1:200 | N/A | Abcam | ab181469 | Alexa Fluor 488 | 1:400 | Thermo Fisher | A11001 |
| **pSTAT3** | 1:100 | N/A | Cell Signaling | 4113T | Alexa Fluor 488 | 1:400 | Thermo Fisher | A11001 |
| **pJAK2** | 1:100 | Alexa Fluor 647 | Abcam | ab200340 |  |  |  |  |
| **Phalloidin** | 1:1000 | iFluor 647 | Abcam | ab176759 |  |  |  |  |
| **Phalloidin** | 1:1000 | iFluor 555 | Abcam | ab176756 |  |  |  |  |
| **Hoechst 33342** | 1:2000 | N/A | Invitrogen | H3570 |  |  |  |  |

